# Dendritic cells focus CTL responses toward highly conserved and topologically important HIV epitopes

**DOI:** 10.1101/2020.07.13.200527

**Authors:** Tatiana M. Garcia-Bates, Mariana L. Palma, Denise C. Hsu, Jintanat Ananworanich, Bette T. Korber, Gaurav D. Gaiha, Nittaya Phanuphak, Rasmi Thomas, Sodsai Tovanabutra, Bruce D. Walker, John W. Mellors, Paolo A. Piazza, Eugene Kroon, Sharon A. Riddler, Nelson L. Michael, Charles R. Rinaldo, Robbie B. Mailliard, on behalf of the I4C and RV254 Study Groups

## Abstract

During early HIV Infection, immunodominant T cell responses to highly variable epitopes lead to the selection and expansion of immune escape variants. As a potential therapeutic strategy, we assessed a specialized type 1-polarized monocyte-derived DC dendritic cell (MDC1)-based approach to selectively elicit functional CD8^+^ cytotoxic T lymphocyte (CTL) responses against highly conserved and topologically important HIV epitopes. Cells were obtained from 10 HIV-infected individuals in the Thailand RV254/SEARH010 cohort who initiated suppressive anti-retroviral therapy (ART) during Fiebig stages I to IV of early infection. Autologous MDC1 were generated for use as peptide antigen presenting cells to induce *ex vivo* CTL responses against HIV Gag, Pol, Env and Nef. Ultra-conserved (Epigraph) or topologically important (Network) antigens were respectively identified using the Epigraph tool and a structure-based network analysis approach, and each compared to overlapping peptides spanning the entire Gag proteome. MDC1 loaded with either overlapping Gag, Epigraph, or Network 14-21mer peptide pools were consistently capable of activating and expanding HIV-specific T cells to epitopes identified at the 9-13mer peptide level. Some CTL responses occurred outside of known or expected HLA associations, providing evidence of new HLA-associated CTL epitopes. Comparative analyses of peptide pools demonstrated more sequence conservation among the Epigraph antigens, but statistically higher magnitude of CTL responses to Network and Gag peptide groups. Importantly, when select Gag antigens used to initiate the cultures were part of the Network peptide pool, CTL responses directed against these topologically important epitopes were enhanced as compared to when they were included within the complete pool of overlapping Gag peptides. Our study supports that MDC1 can be used to effectively focus CTL responses toward potentially fitness-constrained regions of HIV as a therapeutic strategy to prevent HIV immune escape and control viral replication.

**Author summary:** A major hurdle in the development of a successful HIV immunotherapy is the capacity of the virus to evade the immune response by efficiently establishing epitope variants in response to selective pressure. While effective at suppressing viremia, current regimens of antiretroviral therapy (ART) are not curative. Therefore, achieving immune control of HIV upon cessation of ART as a functional cure, similar to that observed in ‘elite controllers’ (EC), has been a major therapeutic goal. Such immune control is realized through the actions of antigen-specific cytotoxic T cell lymphocytes (CTL) capable of specifically targeting sequence-conserved epitopes in HIV. In this study, a specialized, antigen presenting, dendritic cell (DC)-based vaccine strategy was used to elicit HIV specific CTL responses *in vitro* against carefully selected, ultra-conserved and topologically important epitopes. This DC-based approach yielded broad responses against peptide epitopes of both known and unknown HLA-associations, the latter of which implies the uncovering of potentially novel epitopes. Importantly, we demonstrate that CTL responses can be re-directed or focused toward potentially more fitness-constrained regions of the virus, thus highlighting the potential for DC-based therapies to induce immune responses that circumvent the issue of viral escape.

## Introduction

Adaptive immune pressure and viral fitness restrictions in untreated HIV infection result in distinct regions of low and high diversity in the viral genome, with the low diversity regions being a preferred antigenic target of immunotherapy [1]. Beginning during acute HIV infection (AHI), immunodominant T cell responses tend to be towards highly variable viral epitopes that can rapidly lead to immune escape variants [2, 3]. However, the antigenic diversity of the HIV population and establishment of CTL escape variants within an individual are less in persons who initiate antiretroviral therapy (ART) during early stages of HIV infection compared to those initiating ART during the progressive, chronic infection [4]. Therefore, implementing a ‘shock and kill’ or ‘kick and kill’ immunotherapeutic approach [5] in those individuals who begin ART during earlier stages of infection could effectively target latently infected CD4^+^ T cells harboring replication-competent HIV.

A major challenge to assessing the ‘kick and kill’ hypothesis as a strategy for immunotherapy of HIV infection is a safe and efficient approach to elicit functional CD8^+^ cytotoxic T lymphocyte (CTL) responses to fitness-constrained viral epitopes. Our strategy for immunotherapy of HIV infection centers on myeloid dendritic cells (DC), professional antigen presenting cells (APC) that induce potent antigen-specific T cell responses in immunotherapy trials for end-stage cancers [6–8]. An advantage of this strategy is that it enables effective presentation of very short regions of protein. This facilitated us to focus the immune response on regions containing epitopes that are either extremely conserved globally, or of potential value due to spanning topologically important amino acids [9]. This is supported by our evidence that DC induce highly potent memory CD8^+^ T cell responses to HIV against a broad array of MHC class I epitopes *in vitro* [10, 11]. Moreover, DC have been safely used in clinical immunotherapy trials for HIV [12, 13], with the current form of DC immunotherapy resulting in a significant if temporary delay in HIV rebound after stopping ART that is related to enhanced T cell control of HIV replication [12, 14].

We hypothesize that the DC used in HIV immunotherapy trials to date have not been adequately equipped with the characteristics needed to specifically direct and support effective type 1-biased cellular immune responses that are required to successfully combat cancers and intracellular infections such as HIV [10, 15–18]. In fact, the methods commonly used to generate mature DC, including the use of maturation factors such as prostaglandin E2 (PGE2) and CD40L, typically give rise to mature DC that quickly become deficient in their capacity to produce IL-12p70 [19], a critical Th1 and CTL driving factor [20]. Indeed, we have found that naïve CD8^+^ T cells from individuals with chronic HIV infection [21] and uninfected individuals [22] can be primed *ex vivo* with autologous, high IL-12p70 producing, type 1-polarized monocyte-derived DC (MDC1) to become efficient cytotoxic T lymphocytes (CTL). Moreover, pre-existing memory CD8^+^ T cells present during chronic HIV infection are capable of recognizing CD4^+^ T cell targets expressing established variant HIV epitopes to produce inflammatory cytokines, but are predominantly dysfunctional in their killing capacity [21–24] and display signatures of immune exhaustion [25]. However, these newly MDC1-primed CTL are effective at killing CD4^+^ T cells infected with autologous HIV [21].

We propose that an effective, functional HIV cure will need to overcome the emergence of early CTL escape variants and immune exhaustion by centering on a carefully designed, MDC1-based immunotherapy targeting highly conserved epitopes or topologically important regions of the HIV proteome that are structurally and functionally critical to viral fitness [9, 26]. Our overarching hypothesis is that MDC1 can be an immunotherapeutic tool to effectively correct or focus CTL activity toward highly conserved or topologically important HIV antigenic sites in those who begin ART during AHI. We therefore tested this approach utilizing a subset of participants in the well-defined RV254 cohort in Thailand who initiated ART during early HIV infection (Fiebig stages I-IV) [27].

We applied two diverse but complementary methods to select CTL antigenic targets. The first method we used was the graph-theory based, computational approach, Epigraph90 [26, 28, 29], which enable us to identify conserved HIV peptide libraries to optimize vaccine coverage of potential CD8^+^ T cell epitope (PTE) variants found in the diverse HIV population. The algorithm allows for exploration of epitope features relevant to an immunotherapeutic DC vaccine design that were previously intractable, such as balancing the costs in PTE coverage with rare epitope exclusion and optimizing coverage of *in vivo* diversity. The Epigraph approach was thus used to define short regions (14-21 amino acids in length) of the proteome with extremely high conservation levels at the global population level; the included regions contained multiple known and/or predicted CD8^+^ T cell epitopes and conserved regions for with-in subject targeting [26]. The focus on extremely conserved but short peptide fragments is particularly well suited to MDC1-priming for CTL induction [22], but similar vaccine antigen design strategies have shown that the immune response can be refocused towards highly conserved elements using DNA delivery [30, 31]. Also, longer regions of the proteome that contain *relatively* conserved regions (balancing the inclusion of more potential epitopes with less stringent conservation requirements) can also help focus the immune response on more conserved regions that are beneficial in terms of clinical outcomes [32–34], using vector delivery strategies or self-amplifying mRNA [32, 35].

The second method for selecting peptide antigens employed a structure-based network analysis to identify structurally and functionally constrained epitopes [9]. Structural data were used to build networks of non-covalent interactions between amino acid side chains, and subsequently analyzed by graph theory metrics to quantify the sum contribution of each residue to the protein’s global architecture. The scientific premise and rationale of this network theory is to identify amino acid residues of topological importance, which are critical to a protein’s structure and function [9]. Thus, effective immune targeting of these highly networked regions of the viral proteome would greatly and negatively impact viral fitness.

## Results

### Characteristics of the RV254/SEARCH010 study cohort

The specimens used in our study were from a well-characterized RV254/SEARCH010 study cohort of adults who were diagnosed with AHI based on HIV screening at the Thai Red Cross Anonymous Clinic as described in the materials and methods [36]. The participants in our study were all men who started virus-suppressive ART during Fiebig I (n=2), Fiebig II (n= 2), Fiebig III (n=4), or Fiebig IV (n=2) stages of early HIV infection based on the Fiebig staging system [37] (Table 1). The HLA alleles of the respective participants are listed in Table 1. Samples used in our experiments were from blood specimens collected between weeks 48 to 240 post-ART initiation, from which PBMC were isolated and stored for future use. Plasma viremia loads at these time points were all bellow 20-50 copies per milliliter. The median CD4^+^ T cell count was 559 cells/ml (IQR 465-687), and the CD8^+^ T cell count median was 496 (IQR 435-596) (Table 1).

**Table 1.**
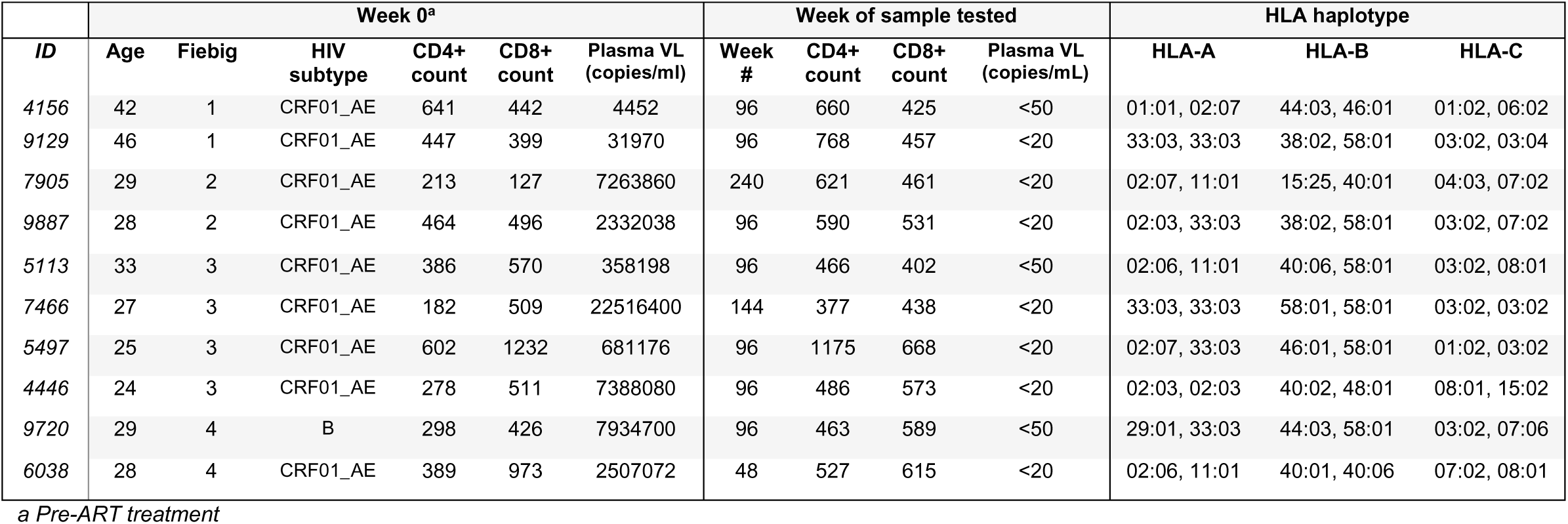
Demographic Characteristics of RV254 Cohort.

### Distinct methods used to identify highly conserved and topologically important CTL antigenic targets

We used 3 distinct approaches for selecting the different sets of peptide immunogens for our study. The first set consisted of a full-length HIV Gag protein peptide pool (referred to as “Gag” for simplicity) comprised of both conserved and non-conserved regions of Gag, which served as a reference point for our study because Gag specific responses are associated with viral control in natural infection, including those in highly variable p17 protein [38](Fig 1). In order to refine immunogen design and to select peptides representing conserved regions of HIV Gag, Env and Pol, two approaches were used as described in the materials and methods. We defined one method as the “Network” based design, which is founded on a structure-based network analysis that identifies topologically important epitopes within a protein [9]. The second method was defined as “Epigraph”, which is based on a highly efficient algorithm that can be used to define conserved HIV peptides as potential CD8^+^ T cell epitopes (PTE) and to define complementary set of antigens that can provide optimal population coverage of potential epitopes across diverse viruses [26, 28, 29]. The 3 peptide pools that were tested in our antigenicity studies consisted of peptides ranging from 14 to 21 amino acids in length (14-21mer, Fig 1). The full-length Gag peptide pool consisted of 45 peptides that overlapped by 10 amino acids and spanned Gag p17, p24, p7 and p6 proteins of the HIV-1 subtype CRF01-AE, the predominant strain in Asia. Of these 45 peptides, 44 were 21mers and 1 was a 14mer. Network peptides consisted of 25 15-21mer peptides that were combinations of Env (n= 4), Nef (n=2), Pol (n=12) and Gag (n=7). Epigraph peptides consisted of 40 14-21mers comprised of Gag (n=5) and Pol (n=35). It is important to note that the peptide lengths for the three groups differed, with the Gag group having 98% 21mer vs. 2% 14-19mer, the Network group with 92% 21mer vs. 8% 15-17mer, and the Epigraph group with 47.5% 21mer vs. 52.5% 14-19mer. For *in vitro* T cell stimulation studies, monocyte-derived MDC1 were loaded with the pooled peptides and used as APC for inducing autologous CD4^+^ and CD8^+^ T cell responses.

**Fig 1.**
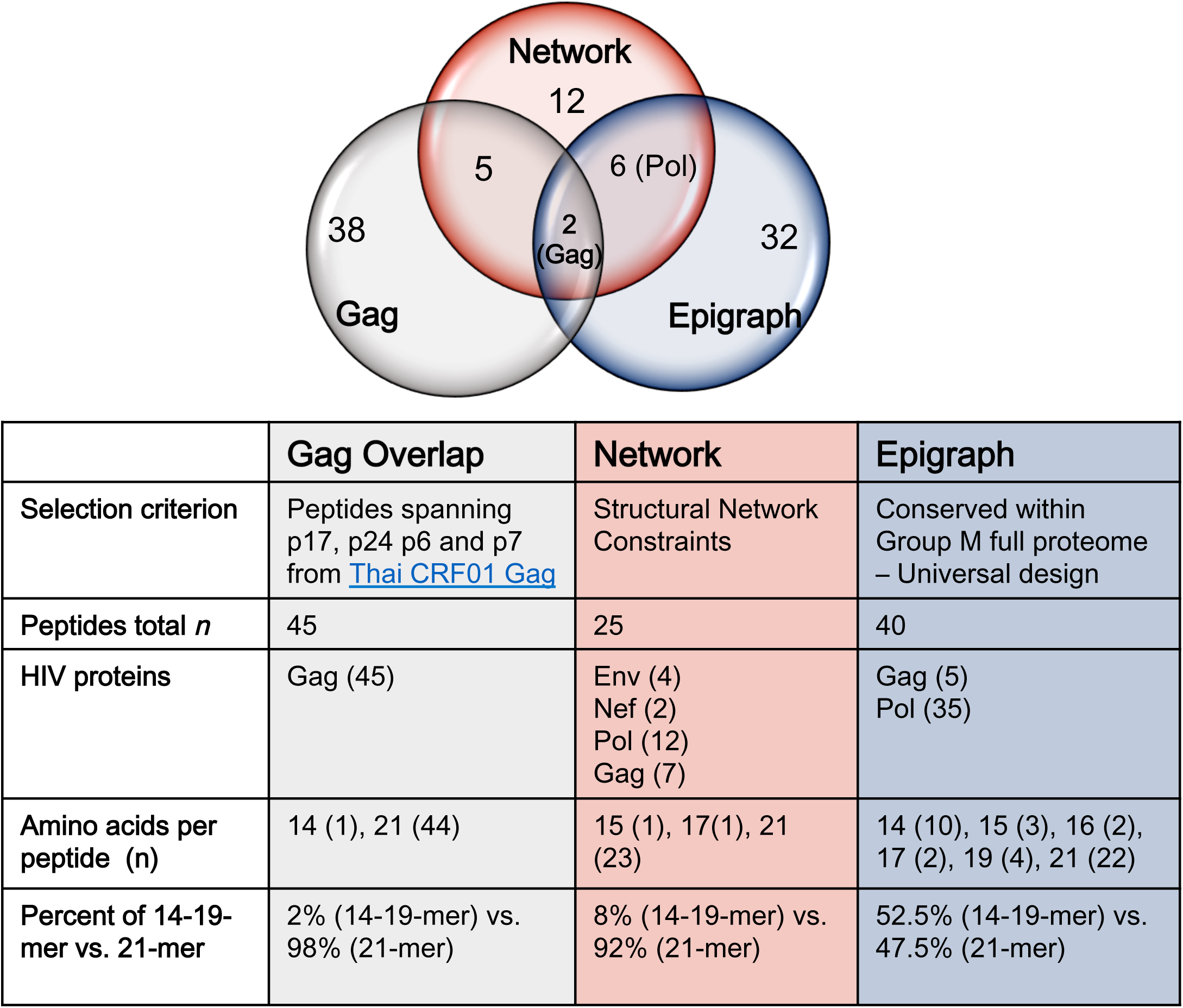
Description of the three peptide pools used in the study. The full-length HIV Gag protein peptide pool (Gag Overlap) comprised of both conserved and non-conserved regions of Gag. Two distinct methods were used to identify HIV peptides for topologically important and highly conserved CTL antigenic targets (Network and Epigraph) are described in materials and methods. Selection of conserved regions were initially focused in the 5’ half of the genome (Gag and Pol) using the Epigraph method, as to facilitate downstream sequencing of clinical samples. However, this constraint was lifted when using the Network method and some conserved regions within Env and Nef (3’ half of the genome) were added to this peptide pool.

### Unrefined evaluation of efferent HIV-1 specific T cell responses initially induced by antigen presenting autologous MDC1

To test for induction and long-term expansion of antigen-specific T cells responsive to the peptide pools described above, we performed a 21 day *in vitro* stimulation assay using antigen-presenting, mature, monocyte-derived DC that were polarized toward a high IL-12p70-producing type-1 phenotype (MDC1) as described previously [39]. The MDC1 were either left untreated (Empty) or loaded with the pool of Gag peptides (Gag), Network peptides (Network), or Epigraph peptides (Epigraph), and subsequently used for *in vitro* stimulation and expansion of isolated autologous T cell responders in long-term co-cultures. Using the same peptide antigen pool used to initiate the DC:T cell co-cultures, the expanded T cells were tested for their respective recall responsiveness to secondary antigenic stimulation by IFNγ ELISpot assay and flow cytometry analysis (see materials and methods) (Fig 2A).

**Fig 2.**
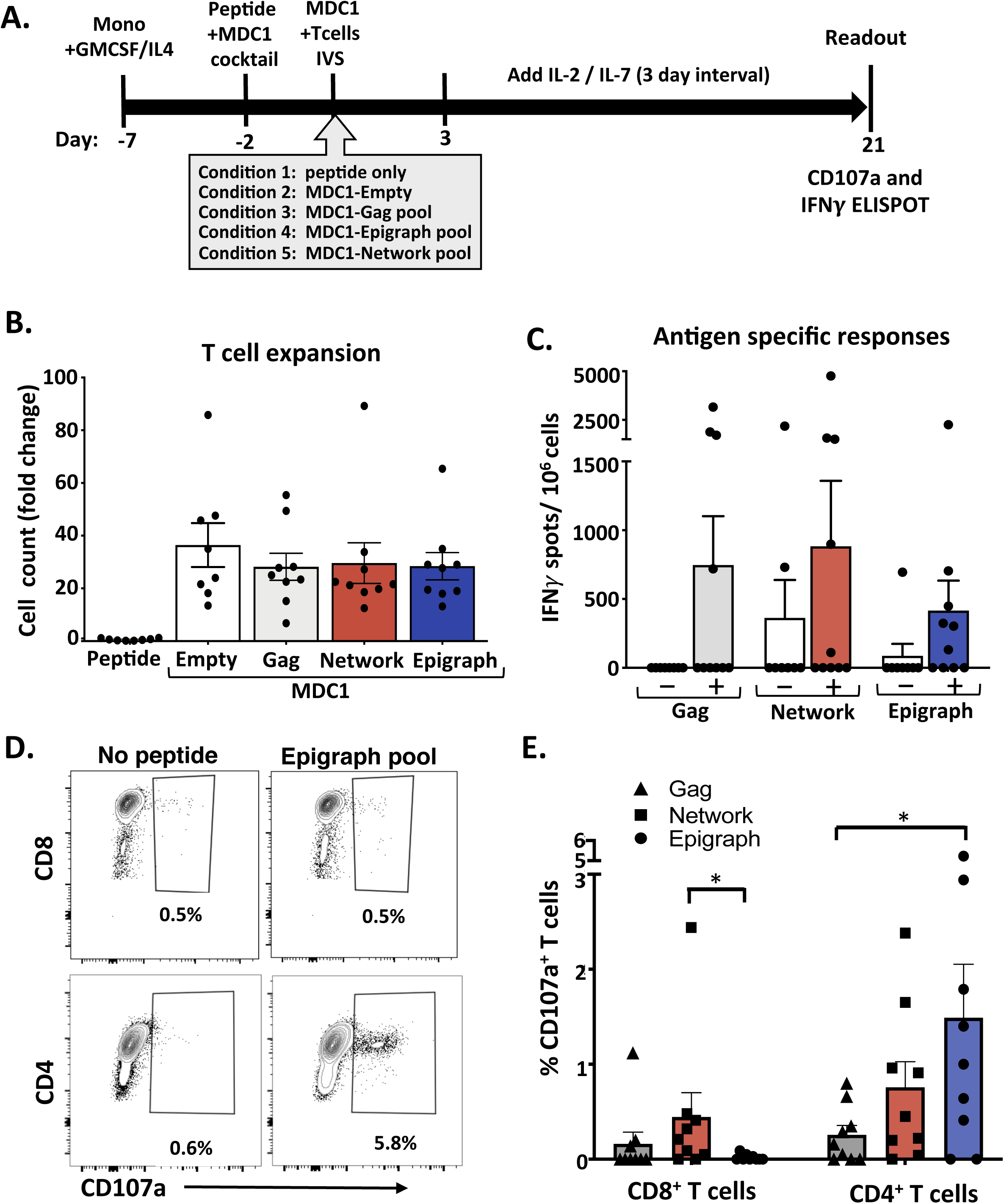
Unrefined evaluation of efferent HIV-1 specific T cell responses initially induced by antigen presenting autologous dendritic cells. A) Timeline of experimental conditions where monocytes isolated from PBMC (Day -7) and were treated with GMCSF and IL-4. After 5 days (Day -2) the iDC were treated with the MDC1 Th1-polarizing cocktail and exposed to either DMSO (Empty, Condition 2) or to one of 3 peptide pools (Gag, Network and Epigraph), shown as Conditions 3, 4 and 5 respectively. T cells stimulated with peptide pool only (without DC) served as an additional control (Condition 1). After 48h (Day 0) the differentially treated MDC1 were co were cocultured with autologous T cells. After 21 days, the T cells were assessed for net expansion and antigen specific responses to a secondary exposure to the respective peptide pools. B) T cell expansion was determined by counting the *in vitro* sensitized T cells at day 21. Results are shown as fold change above the number of T cells used to initiate the cultures at day 0. C) T cell cultures that were expanded in the presence of MDC1 loaded with the Gag, Network or Epigraph peptide pools (respectively represented by + symbols) were tested for induced IFNγ responses to secondary exposure to the respective Gag, Network, or Epigraph (Y axis) peptide pools by ELISpot assay. T cells non-specifically expanded by the ‘Empty’ control MDC1 (-) were also tested for responsiveness to each of the respective peptide pools. Results are shown as spot forming units per million cells (IFNγ SFU/10^6^ cells). D) Representative flow cytometry plots of one participant illustrating the gating on Epigraph peptide pool responding CD107a^+^ CD8^+^ T cells (top panels) and CD107a^+^CD4^+^ T cells (bottom panels). E) Graphical representation of the percent of specific peptide induced CD107a^+^ expressing CD8^+^ T cells (left) and CD4^+^ T cells (right) in 9 study participants tested. *p<0.05

While T cell expansion occurred equally in all of the culture conditions that contained MDC1, even in the absence of exogenous peptide (Empty), the T cells failed to expand in culture in the absence of MDC1 (Fig 2B). Antigen-induced IFNγ ELISpot responses were detected in all of the peptide groups tested (Gag, Network, Epigraph), with no significant differences in the cumulative magnitude of responses noted among these pools, although it is important to note that there was a different number of epitopes incorporated into each peptide pool (Gag > Epigraph > Network) (Fig 2C). Importantly, T cell cultures that were non-specifically expanded using non-peptide loaded, control (Empty) MDC1 yielded few antigen responsive cells, with the exception of 2 participants whose cells responded to the Network peptides during the assay readout, highlighting the importance of both MDC1 and peptide antigen for the selective induction and long-term survival of HIV antigen-specific T cells. Since the expanded cultures included both CD4^+^ and CD8^+^ T cells, the ELISpot assay could not distinguish the relative contribution of the responses made by each T cell subset. Therefore, to differentiate between CD4^+^ and CD8^+^ T cell responses, we used flow cytometry analysis to test the relative responsiveness of these individual T cell subsets to the HIV peptides based on their induced expression of CD107a after a 6 h stimulation with their respective peptide pool (Fig 2D, E). We found a higher percentage of antigen-responsive CD4^+^ T cells in all groups tested compared to CD8^+^ T cells. This is demonstrated in Fig 2D, with representative flow cytometry plots of one study participant’s responses to the entire Epigraph peptide pool, where the CD4^+^ T cell response reached 5.8% compared to a 0.5% response in the CD8^+^ T cell fraction. In particular, the percent of antigen responsive CD8^+^CD107a^+^ T cells was lowest for those cultures generated using MDC1 loaded with either the Gag or Epigraph peptide pools, with a mean of 0.16% (range 0% to 1.1%) and 0.02% (range 0% to 0.09%) respectively. Cultures generated using the MDC1 loaded with the conserved Network peptides also yielded relatively low CD8^+^ T cell responses, although the overall percentage of antigen responsive CD8^+^CD107a^+^ T cells in this group was significantly higher than the others, with a mean of 0.45% (range 0% to 2.4%) (Fig 2E). Interestingly and in contrast to our results with the CD8^+^ T cells, the highest percentage of antigen-responsive CD4^+^CD107a^+^ T cells was generated using MDC1 loaded with the Epigraph peptide pool (mean 1.5%, range 0% to 5.2%), which was significantly higher than those induced using MDC1 loaded with the Gag peptide pool (mean 0.26%, range 0 to 0.8%), and higher (although not statistically significant) than the Network peptide pool (mean 0.76%, range 0 to 2.4%).

### Efferent CD8^+^ T cell responses become evident with refined analysis using 9-13mer peptide epitopes

We hypothesized that the observed overall higher responses found among CD4^+^ T cells compared to CD8^+^ T cells were due to the use of longer peptide antigens as direct stimulators in these short-term efferent readout assays. In accordance with previous findings [40], we reasoned that the longer peptides were more readily presented in the context of MHC class II as compared to MHC class I, thus reflecting an inefficient stimulation and detection of the antigen-specific CD8^+^ T cells in the short-term assays rather than the lack of their presence in the expanded T cell cultures. Therefore, we next evaluated whether the efferent CD8^+^ T cell responses induced by direct peptide antigen stimulation was more efficiently revealed by using smaller 9-13mer peptide epitopes that were derived from and contained within the longer peptide sequences used in the initial afferent stimulation by the antigen-presenting MDC1. To demonstrate this, we first generated T cells from a representative HLA A2^+^ study participant using autologous MDC1 loaded with one of the 21mer Gag (Gag144-164) peptides included in the Gag peptide pool, which contained a known HLA-A2-restricted 9mer epitope TV9 (Gag151-159) (Fig 3A). We used the same MDC1-based afferent stimulation strategy as described before, expanding the T cell cultures for 21d, and testing them for secondary efferent response to either the 21mer Gag (Gag144-164) peptide or the 9mer Gag TV9 (Gag151-159) peptide epitope, measuring antigen-induced IFNγ production by ELISPOT assay and by both intracellular cytokine staining (ICS) flow cytometry analysis (Fig 3B and 3C). We observed a mean of 133 SFU/10^6^ cells by ELISPOT (Fig 3B) and 0.08% of the CD8^+^ T cells specifically producing IFNγ when stimulated with the 21mer peptide as determined by flow cytometry analysis (Fig 3C). However, using the 9mer TV9 peptide as the efferent readout-stimulator revealed a much higher percentage of antigen-responsive IFNγ^+^ producing CD8^+^ T cells, with a mean of 1,590 SFU/10^6^ cells by ELISPOT (Fig 3B) and 0.58% being detected by flow cytometry ICS (Fig 3C).

**Fig 3.**
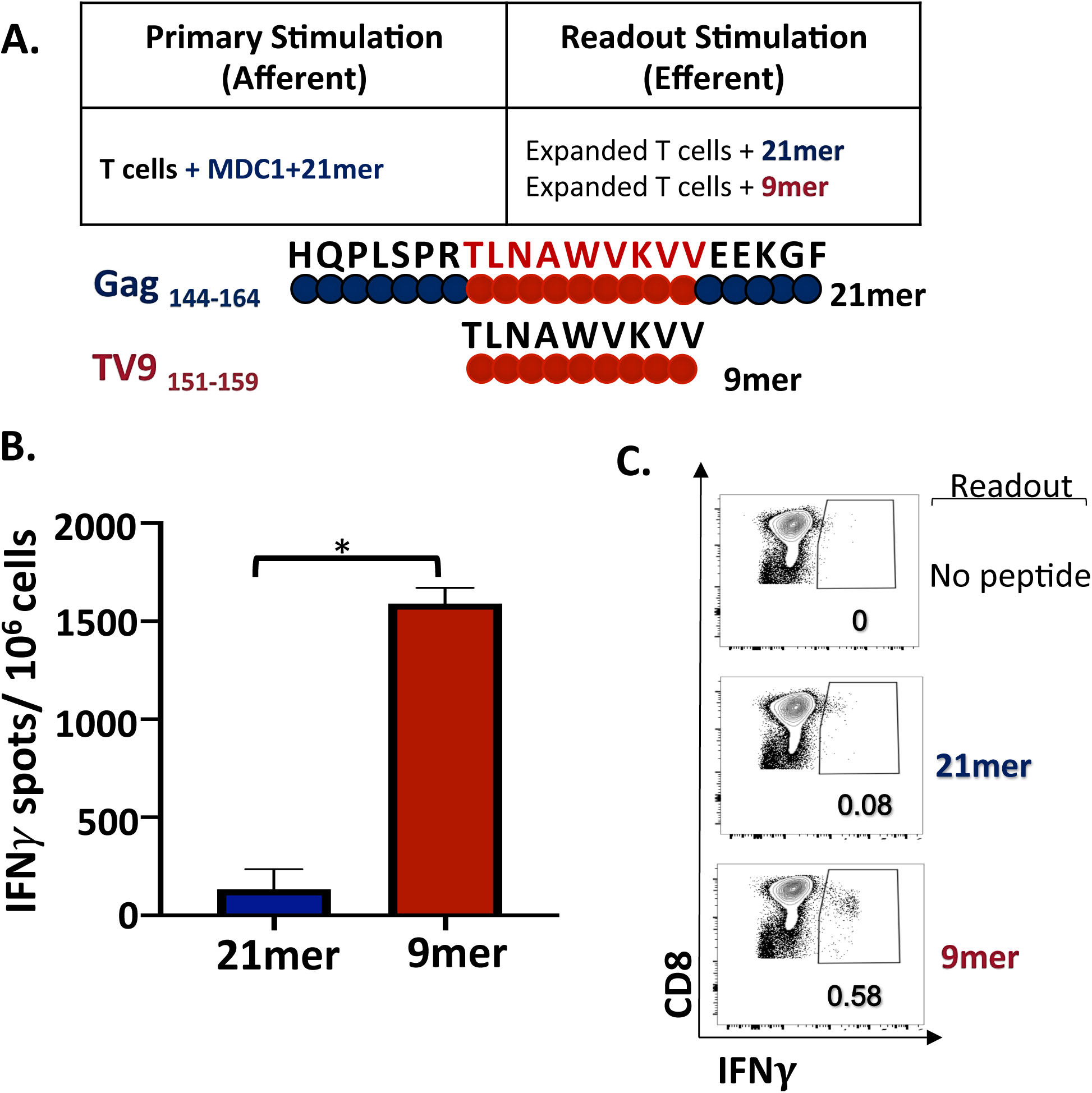
Efferent CD8^+^ T cell responses become evident with refined analysis using 9-13mer peptide epitopes. A) Schematic representation of primary (afferent) and secondary assay readout (efferent) in vitro stimulation conditions of T cell from an HLA-A2^+^ study participant. MDC1 loaded with a 21mer Gag peptide (Gag 144-164) was as the afferent stimulator, and either the same 21mer peptide or the known HLA-A2 restricted 9mer epitope TV9 (Gag 151-159) contained within that 21mer sequence were used as the efferent stimulator in readout assays. B) IFNγ ELISpot assay results showing T cell responses induced by the 9mer and 21mer efferent peptide stimulators, recorded as spot forming units per million cells (IFNγ SFU/10^6^ cells). *p<0.05. C) Antigen peptide-induced IFNγ production by CD8^+^ T cells determined by flow cytometry intracellular cytokine staining (ICS) analysis.

These results prompted us to redesign our CTL readout strategy in order to detect optimal CD8^+^ T cell-specific effector readout responses using smaller peptides derived from the larger 14-21mer peptides used in the initiation of the MDC1:T cell co-cultures. We approached this by selecting shorter 9-13mer sequences using the Los Alamos National Laboratory (LANL) epitope database and the Immune Epitope Database (IEDB), which list known motifs identified as MHC-class I epitopes with the given HLA associations. We narrowed the peptide library selection based on the presence of their sequences within the longer afferent peptides, and their associations with the HLA alleles common to our cohort of Thailand study participants (Tables 2, 3 and 4). An example of a set of smaller efferent assay readout peptides that were selected and derived from one of the larger Gag-associated Network afferent stimulator peptide is described in Fig 4A. In addition, we determined the degree that these sequences matched those shared among the entire HIV-1 M group as well as to the Thailand dominant CRF01 clade specifically, an example of which is shown in Fig 4A and detailed in supplemental figures 2 to 5. A complete list of this analysis for all peptides tested is also included in tables 2, 3, and 4. Of note is that, while the Network associated epitopes individually showed a variable degree of exact matching among the entire M group (S4 Fig), they were more highly matched within the relevant CRF01 clade of this Thai patient population (S5 Fig). Given the algorithm used for their selection, the Epigraph selected peptides were more highly and uniformly matched to both the entire HIV-1 M group (S4 Fig) as well as the Thailand dominant CRF01 clade (S5 Fig). The Gag overlapping peptide group followed a similar pattern to that of the Network group, with a higher degree of variability in exact matching to the entire HIV-1 M group and a relative increase in exact matching to the CRF01 clade.

**Fig 4.**
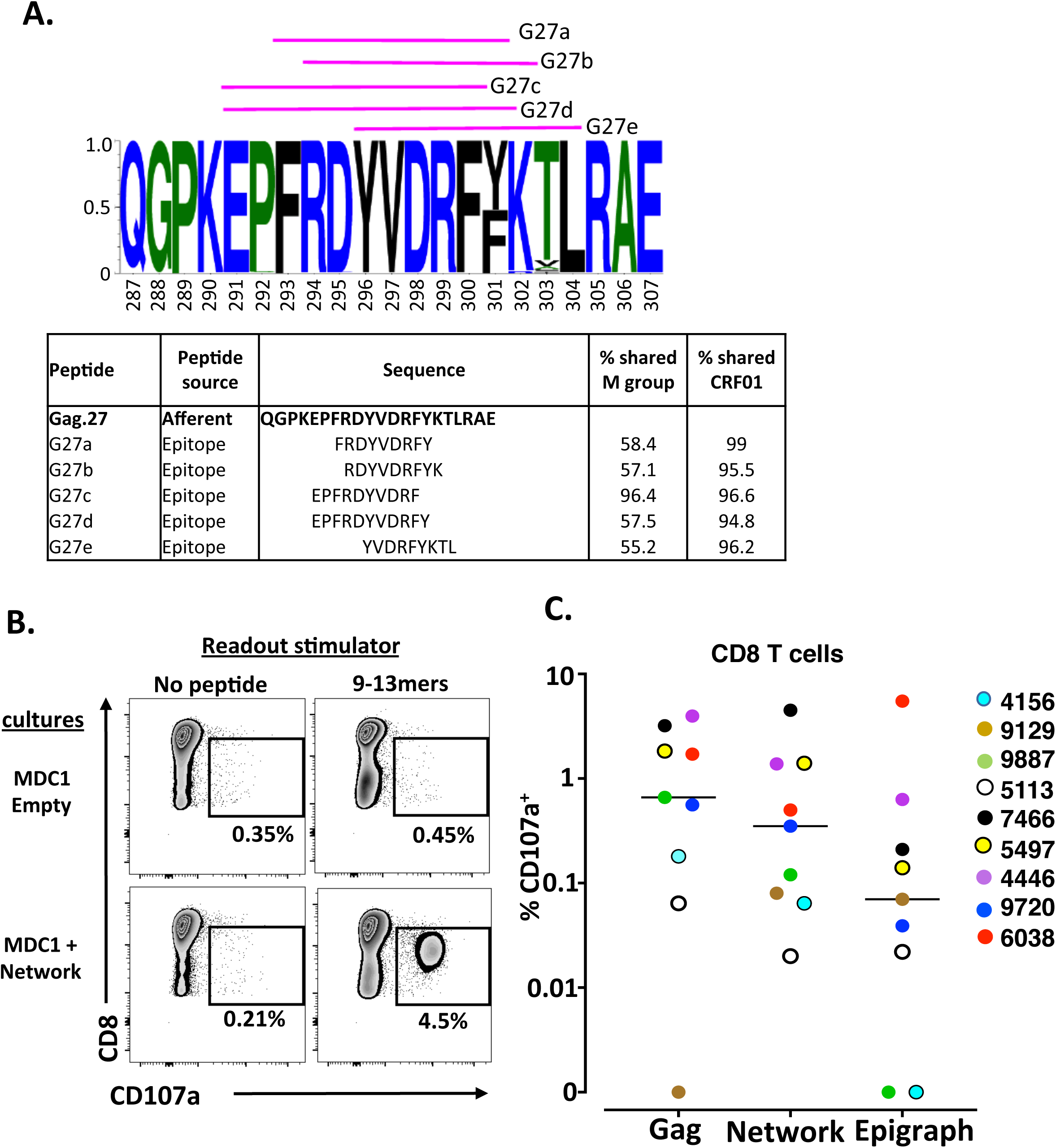
Efferent CD8^+^ T cell responses to 9-13mer HIV antigen peptide pools across all study participants as evaluated by flow cytometry. A) Example of a sequence logo (Gag.27) summarizing the amino acid frequency within the afferent 21-mer peptide. Pink lines represent the shorter 9-mer peptide epitopes used to test efferent responses (also listed in the table). The table lists the peptides and their percentage of exact sequence match with the HIV-1 M group and CRF01 clade. B) Representative flow cytometry data plots generated from one representative study participant illustrating 9-13mer (efferent) peptide antigen-induced expression of CD107a in responding CD8^+^ T cells generated from cultures initiated using MDC1 loaded with the Network (afferent) antigen pool. C) Graphical representation of 9 of each study participant and their percentage of antigen specific CD8^+^ T cell responses generated against the individual Gag, Network and Epigraph peptide pools determined by CD107a expression above background. The lines represent the means of the responses.

**Fig 5.**
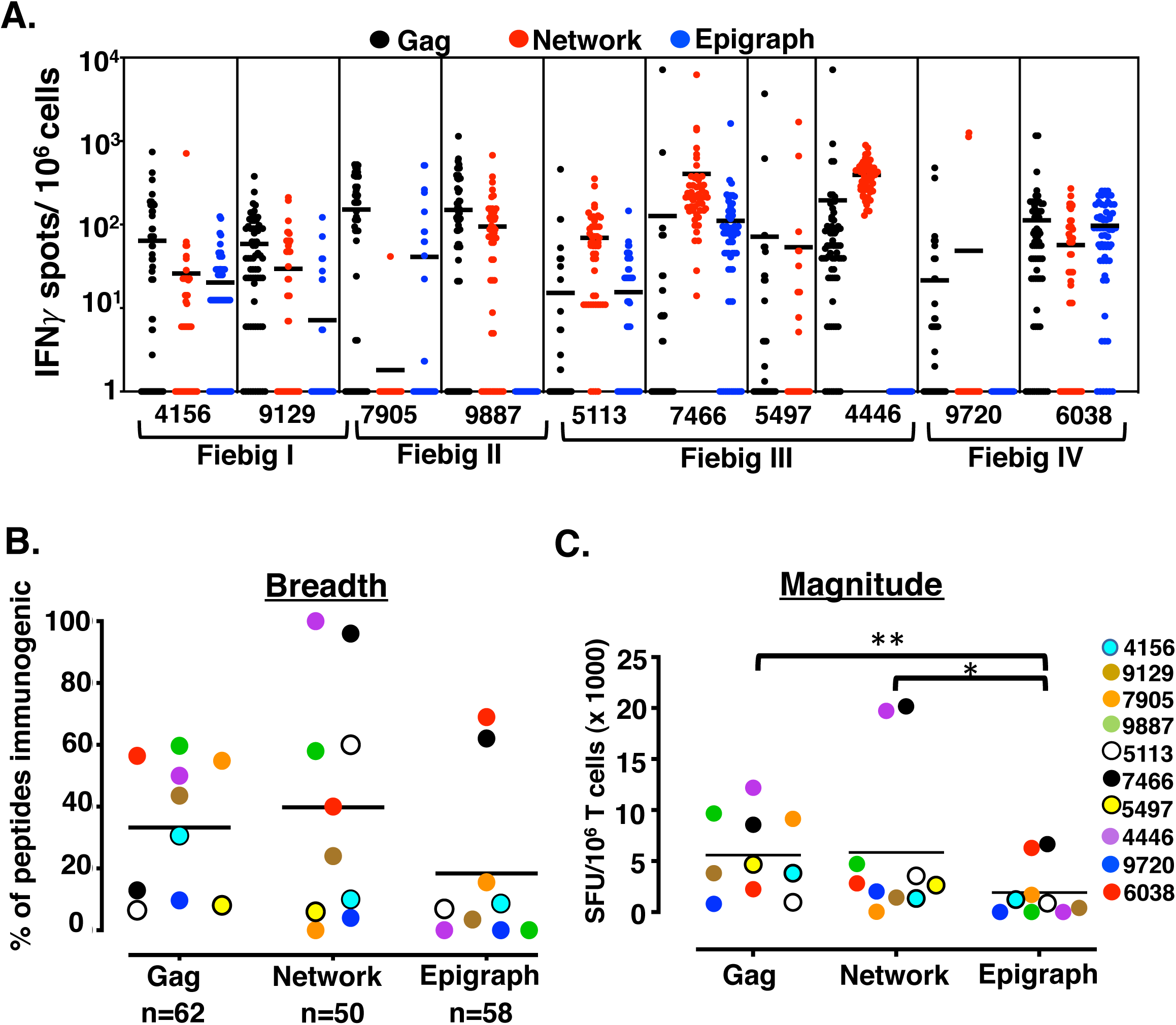
MDC1 induce CTL responses of high heterogeneity against HIV-1 antigenic peptides. A) T cell cultures were initiated in 10 study participants using autologous MDC1 cells loaded with either the Gag, Network or Epigraph pools of 14-21mer peptides. Antigen-induced IFNγ T cell responses generated from each participant were determined at a single peptide level using a matrix of individual 9-13mer peptides derived from the Gag (n=62, black circles), Network (n=50, Red circles) and Epigraph (n=58, blue circles) pools. Results are shown as spot forming units per million cells (IFNγ SFU/10^6^ cells) with each circle representing a response to one peptide. Participant ID# 5497 was not tested for Epigraph peptide responses due to insufficient cell numbers at the onset of the experiment. B) Breadth of the T cell response was quantified by the percent of positive peptide responses (positive responses were >= to 50 SFU/10^6^ cells) within each participant out of the total number of peptides (Gag, n=62; Network, n=50 and Epigraph, n=58). C) Magnitude of the T cell response quantified by compiling the sum of individual peptide-induced responses generated within each peptide group for each participant. *p<0.05, **p<0.01

**Table 2.**
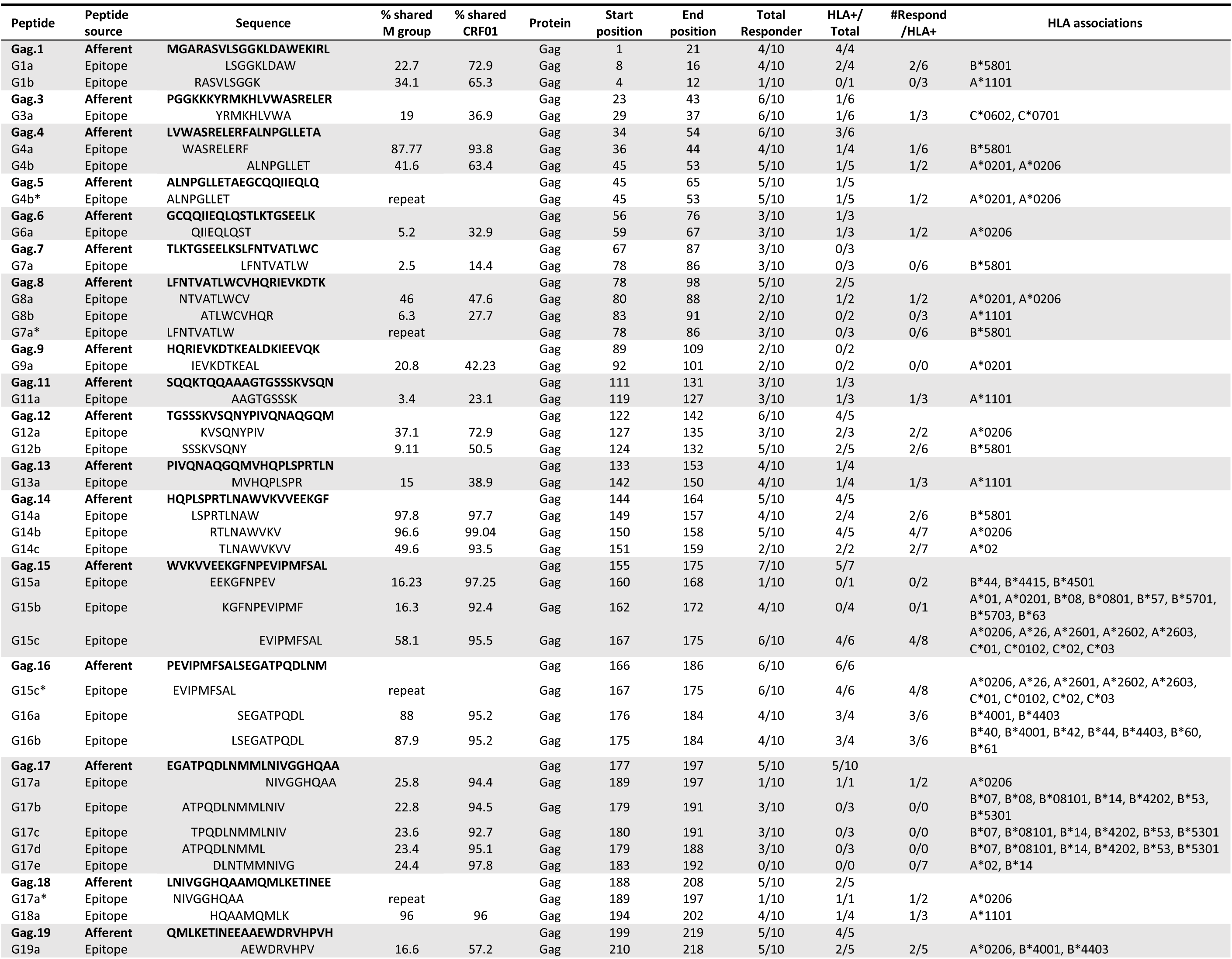
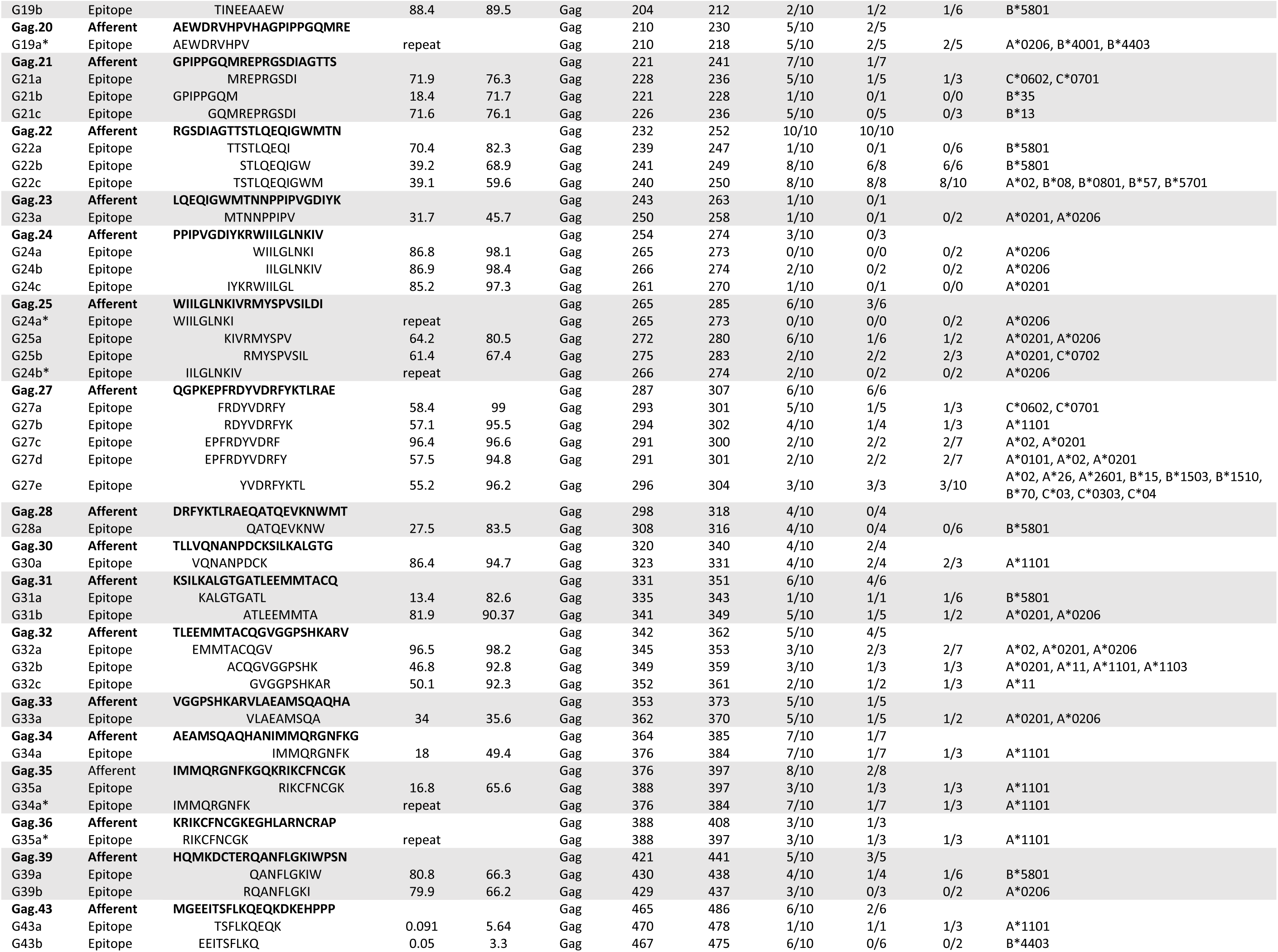
T cell responses to Gag-overlapping pool epitopes.

**Table 3.**
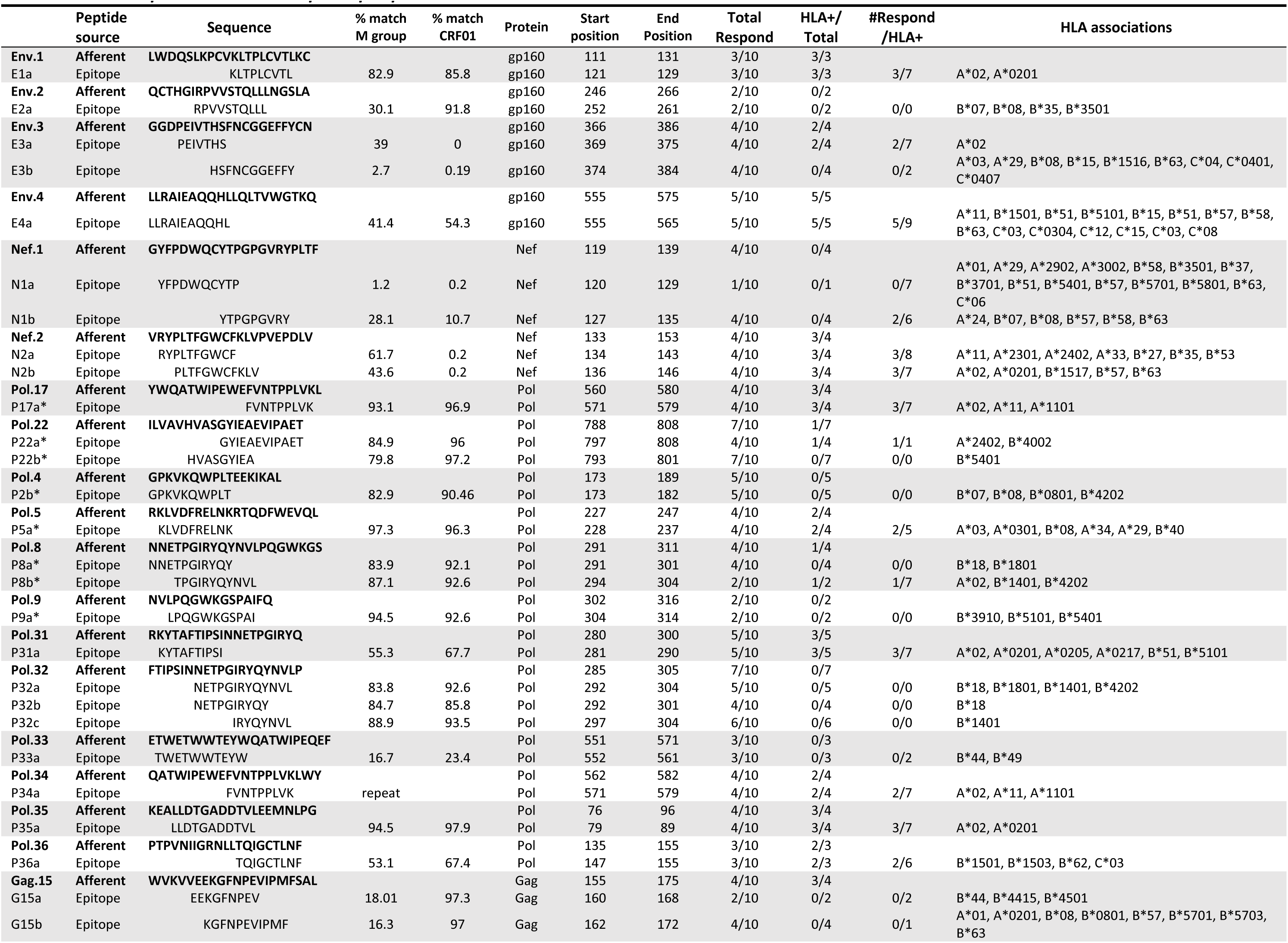
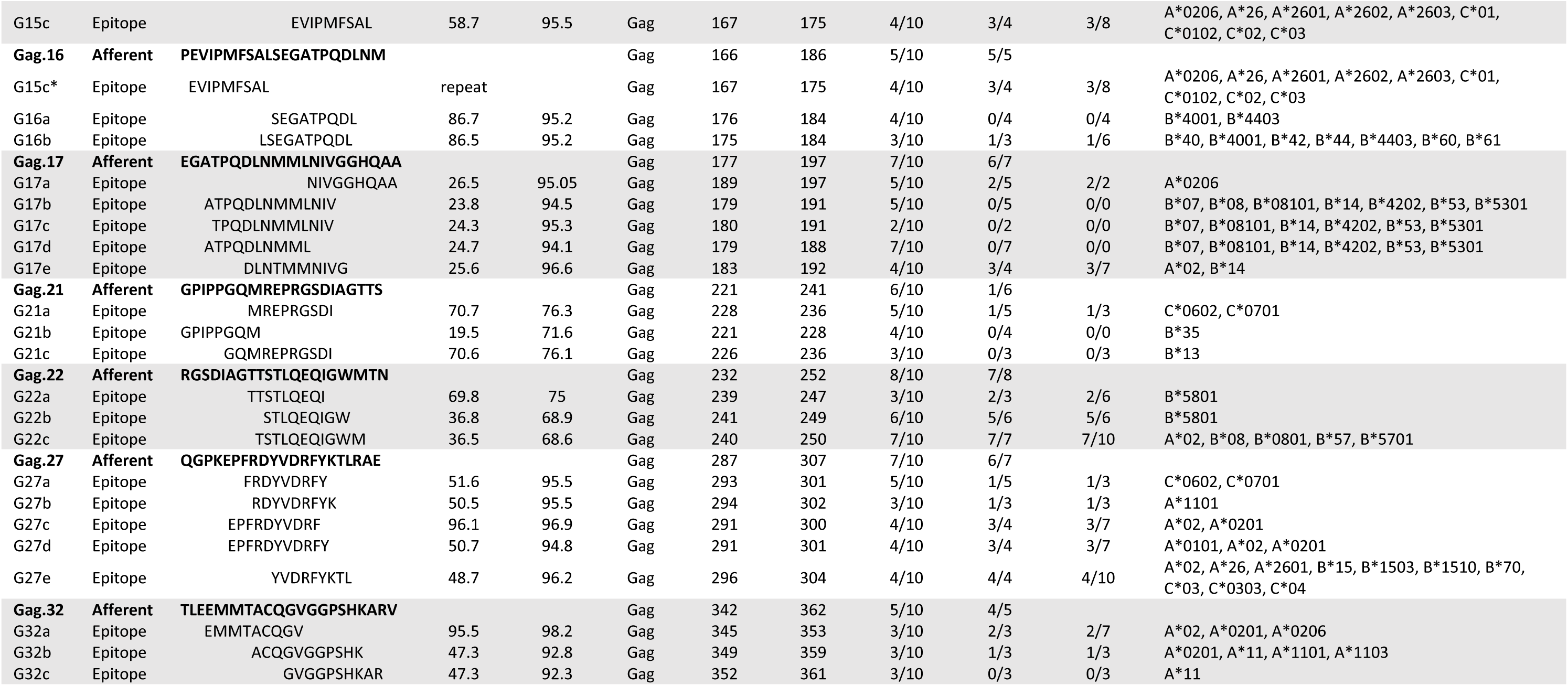
T cell responses to Network pool epitopes.

**Table 4.**
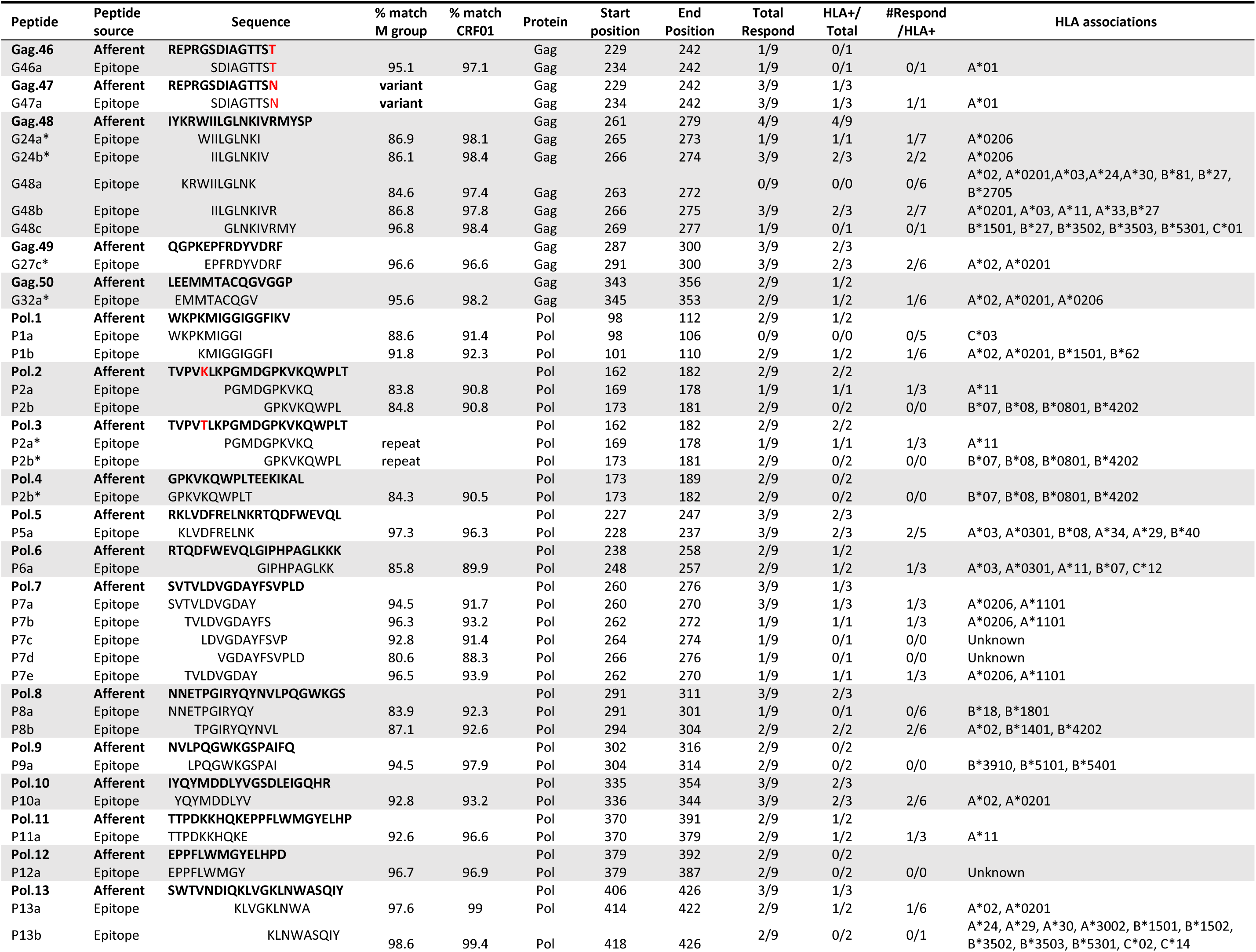
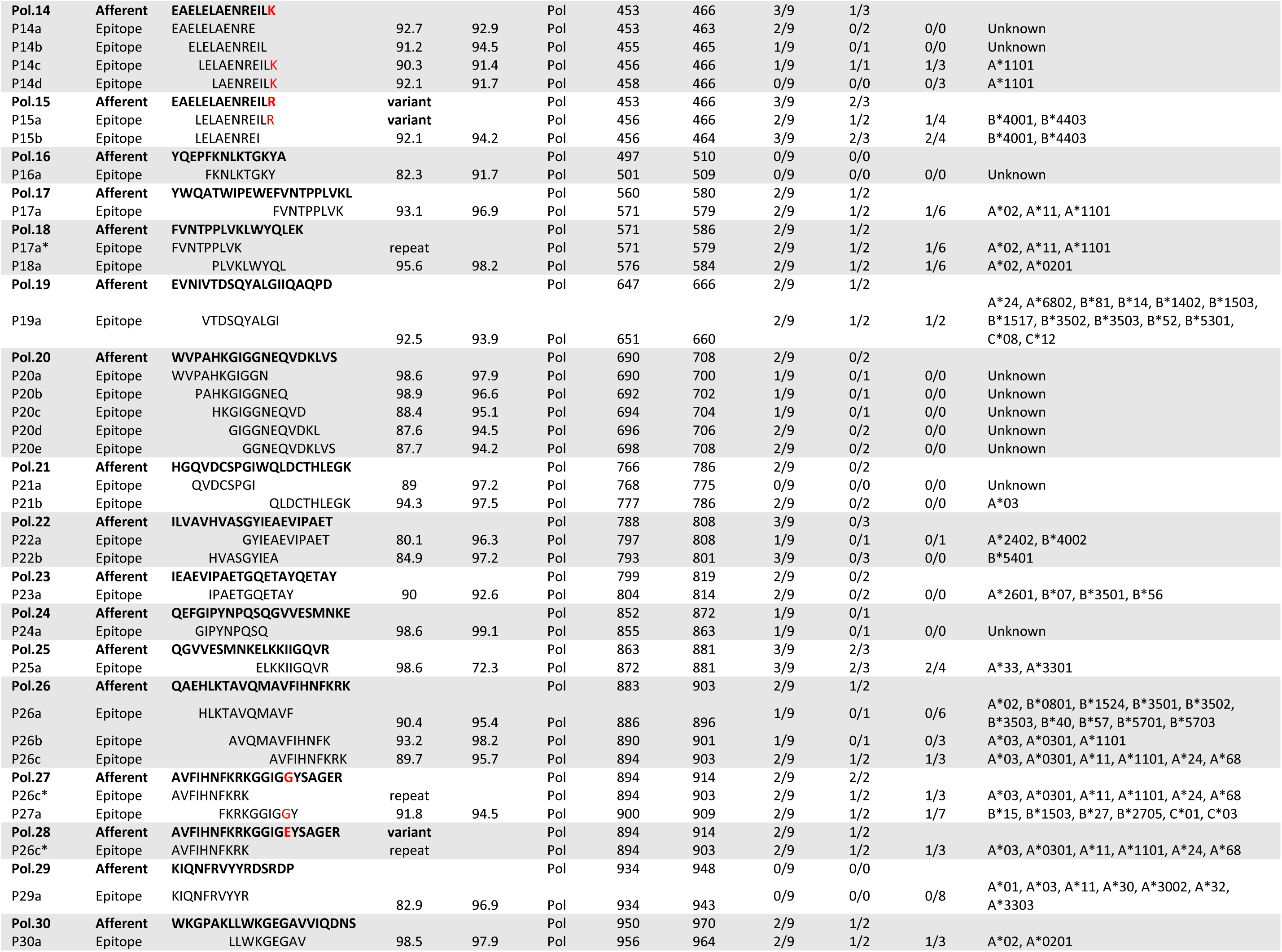
T cell responses to Epigraph pool epitopes.

We repeated the primary *in vitro* stimulation of T cells as described earlier in Fig 2 using MDC1 alone (Empty) or MDC1 loaded with either the Gag, Network or Epigraph peptide pools consisting of the 14-21mers. However, when testing the effector CD8^+^ T cell responses at day 21, this time a pool of relevant 9-13mer peptides served as the efferent stimulator. The results for one representative participant’s efferent, secondary CD8^+^ T cell responses to the Network peptide 9-13mer peptide pool based on the induced expression of CD107a is shown in Fig 4B. We observed a substantial increase in the percentage of antigen-responsive CD107a^+^CD8^+^ T cells revealed using this approach, with 4.5% of the CD8^+^ T cells responding to the Network peptide pool compared to the unstimulated background of 0.21%. Importantly, these responses were not observed in the T cell cultures expanded by MDC1 in the absence of antigen (MDC1-Empty). When we analyzed all participants for antigen-induced CD107a^+^CD8^+^ T cell responses, we found a range of responses to each peptide pool, with most participants reacting to the Gag group (mean 1.35%, median 0.7%), all participants reacting to Network group (mean 0.94%, median 0.35%), and less so to the Epigraph group (mean 0.73%, median 0.07%) (Fig 4C). These data indicate the MDC1 are indeed capable of processing and cross-presenting the larger HIV-1 associated peptides in the context of MHC-class I to induce HIV-1 specific CD8^+^ T cell responses to highly conserved and topologically important regions of the HIV-1 proteome. This also demonstrates that smaller, 9-13mer peptides are required for more accurate quantification antigen-specific CD8^+^ T cell responses.

### MDC1 induce CTL responses of high heterogeneity against HIV-1 antigenic peptides

Each study participant had individual T cell cultures generated using autologous MDC1 stimulator cells loaded with either the larger Gag, Network, or Epigraph peptides. To further analyze the breadth and magnitude of the expanded antigen-specific T cells, the efferent CD8^+^ T cell responses were tested at a single peptide level by evaluating antigen-induced IFNγ secretion by ELISpot using a matrix of 62 individual 9-13mer peptides for the Gag group, 50 peptides for the Network group, and 58 for the Epigraph group. As mentioned earlier, we selected known and predicted CD8^+^ T cell epitopes (by search in the LANL database) contained within the larger 14-21mer sequence that spanned a maximum number of HLA-associations representative of the HLA types of the participants in our cohort. This was done to minimize the number of peptides needed to yield maximum results for each peptide group tested, as the cell number was ultimately a limiting factor. One participant, number 5497, was not tested by the Epigraph pool due to insufficient PBMC availability at the time of initiation of the cultures.

The participants had a broad range of T cell responses to the peptide antigens, with some participants responding to all 3 HIV peptide groups, and others responding to peptides within at least 2 of the groups (Fig 5A). Interestingly, there were particularly high T cell responses generated against several of the 9-13mer peptides among the Fiebig stage III participants, which were especially apparent with the Network peptides. When analyzing participant ID 7466 and 4446 (Fiebig III) in particular, we found these two participants had relatively higher viral loads at week 0, before initiation of ART (Table 1), compared to the other Fiebig III participants. Moreover, we found a positive correlation between viral load at initiation of ART (week 0) and the magnitude of the responses against the Network pool of peptides (S1 Fig). The mean breadth of the CD8^+^ T cell responses to the individual Gag, Network, and Epigraph peptides was 33% (20/62), 40% (20/50), and 18.4% (11/58), respectively (Fig 5B). When analyzing the magnitude of the cumulative responses to each peptide group, both the Gag and Network groups generated significantly higher values than the Epigraph group (Fig 5C).

### Unveiling HLA-associated effector T cell responses to 9-13mer HIV peptides

We analyzed the efferent responses to each of the individual the 9-13mer peptides, and compartmentalized these epitopes based on the respective larger peptides used during the afferent arm of the MDC1-mediated stimulation from which they were derived, as well as to their known or predicted HLA-associations. By doing this, we could predict which of the epitopes was more likely to induce a response based on the individual’s HLA genotype (Fig 6 and Tables 2, 3 and 4). We first quantified the number of individuals that generated antigen-specific effector responses relevant to each 14-21mer peptide used during the initial afferent MDC1-mediated induction of the T cell cultures. Those who had T cells responding to any of relevant 9-13mer epitopes derived from that larger afferent stimulator peptide were determined by IFNγ ELISpot assay, with a value of ≥50 IFNγ SFU/10^6^ cells used as a cutoff for an individual to be considered a responder to that epitope (Fig 6). These results were then matched with the participant’s HLA types (Tables 2,3, and 4).

**Fig 6.**
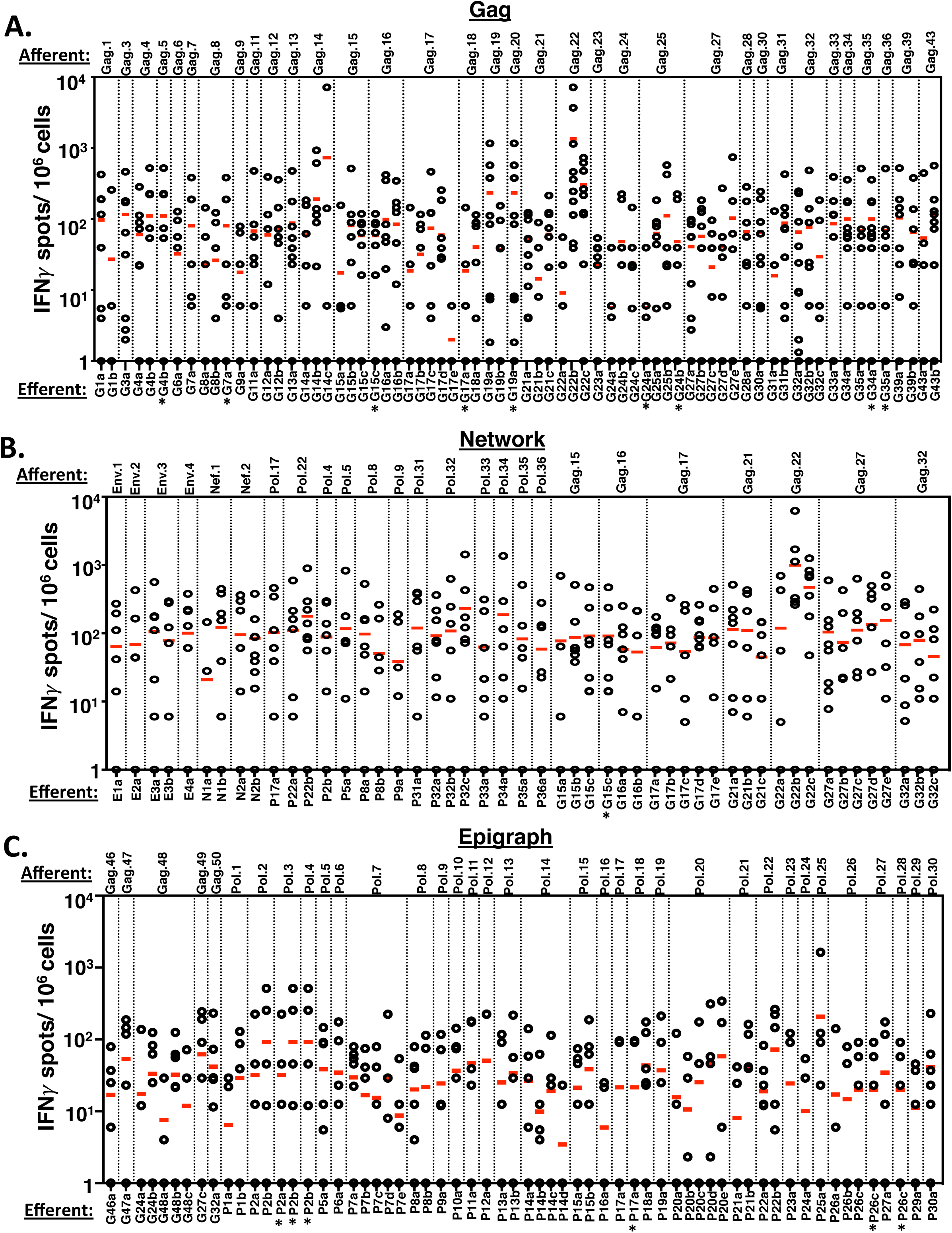
Unveiling 9-13mer peptide HLA restricted T cell responses to HIV antigen pools. T cell responses to 9-13mer single epitopes in the Gag peptide pool (A), Network peptide pool (B), and Epigraph peptide pool (C) were analyzed by IFNγ ELISpot. Responses to individual 9-13 peptides from all 10 study participants were plotted and organized based on the respective larger afferent stimulator peptides used during the initiation of the MDC1: T cell co-cultures. The 14-21mer afferent peptides (top of graphs) and their corresponding efferent assay readout 9-13mer peptides (bottom of graphs) used in the study are listed in Tables 2, 3 and 4. Each plotted circle represents a value generated from 1 of the 10 study participants tested in response to that particular efferent peptide stimulator. A value of ≥50 IFNγ SFU/10^6^ cells was used as a cutoff for an individual to be considered a responder to that epitope.

Of the 34 14-21mer peptides contained in the Gag peptide pool used in the initial MDC1-based T cell stimulation that were assessed, 10 effectively generated cultures yielding antigen-specific effector responses to relevant 9-13mer efferent peptides in at least 5 or more of the 10 study participants tested (Fig 6A, Table 2). One afferent peptide, Gag.22 (sequence RGSDIAGTTSTLQEQIGWMTN), which was present in both the Gag and Network peptide pools, was particularly immunogenic and generated efferent epitope responses in all 10 study participants. Importantly, all of these participants had restricting HLA alleles representative of those capable of binding the Gag.22-associated 9-13mer epitopes. Interestingly, we found that 13 of the larger afferent stimulator peptides from the Gag pool used in the initiation of the T cell cultures yielded specific effector responses to epitopes outside of the known or expected HLA associations of the individual participants (Table 2), suggesting potentially novel epitopes or unreported HLA associations. In the cultures generated using the Network peptide group (Fig 6B and Table 3), we found 11 of 25 of the afferent MDC1 stimulator peptides yielded responses to relevant efferent 9-13mer epitopes in 50% or more of the participants. We also observed 4 afferent peptide antigens that drove efferent peptide responses in 50% or more of the participants, which included responses to peptides outside of expected HLA associations, again indicating the potential discovery of new HLA-associated epitopes. Finally, the Epigraph group also elicited a broad range of responses (Fig 6C and Table 4). While the overall response rate among the participants was not as high as to the Gag and Network peptide groups, with only 1 afferent peptide (Gag.48) from the Epigraph pool approaching a 50% efferent response rate (4 out of 9), 12 of the 35 of the afferent peptides induced responses to their associated 9-13mer peptides in more than 30% of the study participants.

### DC facilitate immune focusing toward subdominant and topologically important epitopes

We hypothesized that HIV-specific T cell responses naturally dominate or become skewed toward immunodominant epitopes in HIV, which can be highly variable, allowing the virus the capacity to easily adapt and escape CTL immune pressure. This can also lead to the establishment of adapted epitopes that drive ineffective cross-reactive memory CTL responses, characterized by their release of cytokines and chemokines in the absence of target killing, thus promoting an inflammatory environment favoring viral dissemination [21–23]. Alternatively, some mutations associated with immune selection pressure impair viral fitness. We posited that MDC1 can be used to facilitate immune focusing of CTL activity toward subdominant HIV epitopes, or to sequences that are most important to maintain protein structures critical to the overall fitness of the virus. Importantly, the Gag peptide pool covered the entire Gag proteome, and it therefore was comprised of both variable and highly conserved Gag-associated epitopes, which included epitopes shared in the select Network peptide pool. This allowed us to directly compare the output (efferent) responses against the same MHC-class I epitopes from T cells derived from cultures that were initiated using MDC1 loaded with either the full-length Gag peptide pool or the select Network peptides. The notion here was to test whether simple elimination of the variable epitopes from the afferent antigen pool used to load the MDC1 stimulators during co-culture initiation would result in enhanced and focused responses toward the conserved and topologically important regions of the virus. Indeed, by limiting exposure of the MDC1 stimulators during the initiation of the T cell cultures to those select Gag peptides contained within the Network pool, as compared to when they are comprised as part of the larger pool of overlapping Gag peptides, resulted in the selective expansion of effector T cells having a significantly enhanced capacity to respond, in both breadth and magnitude, to the same 9-13mer Gag epitopes in the readout assays (Fig 7A). This enhancing effect was noted in total T cell responses generated among all the participants against 21 out of 24 common 9-13mer Gag CTL epitopes tested (Fig 7B). These results demonstrate the utility and potential of using MDC1 to generate and focus effector CTL responses toward conserved and topologically important regions of HIV in those who initiate ART during early HIV-1 infection as a therapeutic strategy to prevent HIV immune escape and control viral replication.

**Fig 7.**
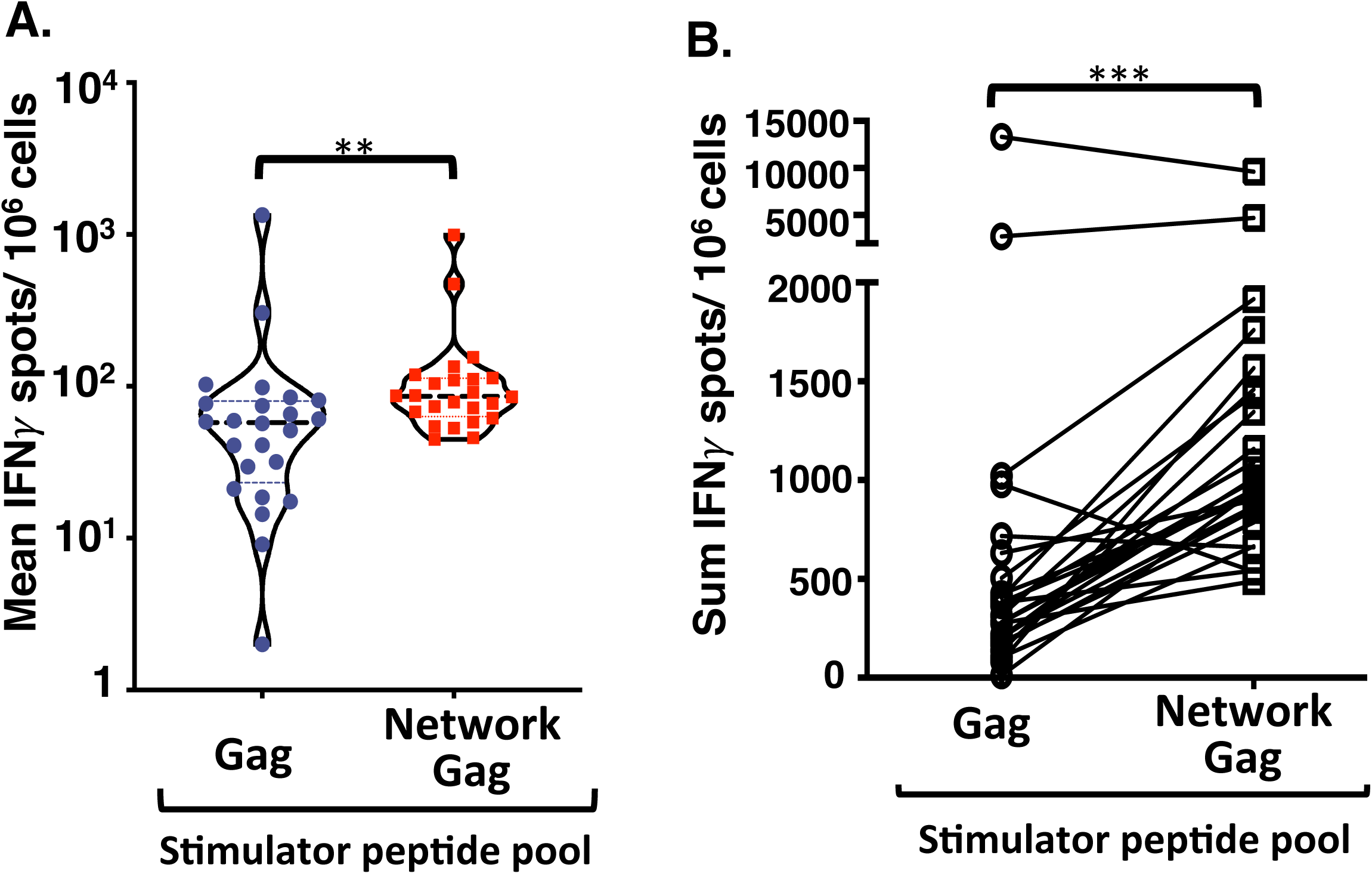
DC facilitate immune focusing toward conserved and topologically important epitopes. MDC1 were loaded with a pool of 14-21mer peptides containing either a mix of overlapping full-length HIV Gag epitopes (Gag) or with the pool of Network peptides also containing Gag associated epitopes (Network Gag), and each were used separately to initiate the activation (afferent) and long-term expansion of responsive T cells. Each dot represents the mean (A) or the sum (B) of the efferent readout responses (IFNγ SFU/10^6^ cells) of each study participant induced against the individual 9-13mer Gag CTL epitopes derived from stimulator peptides common to both afferent stimulator peptide pools.

## Discussion

Major effort has been made toward the design of novel therapeutic strategies to achieve a functional cure for HIV infection, so that HIV-infected individuals would have the capacity to immunologically restrain or inhibit the virus without ART, similar to that observed with HIV EC [41]. Research that has focused on understanding the mechanism of the natural viral control demonstrated by EC has highlighted the importance of generating effective HIV-specific CD8^+^ T cell responses to manage the virus [42, 43]. Importantly, such individuals appear to control viremia by targeting sequence-conserved epitopes derived from topologically important regions of the viral proteome critical to viral fitness [9, 26, 32–34, 44, 45]. Moreover, early ART initiation provides additional benefit for the induction of strong CTL immunity through preservation of CD4^+^ T cell support [46].

In this study, we investigated the induction of HIV-specific T cell responses in early ART treated HIV infected individuals, applying two different analytical approaches to select optimal peptide antigens as potential targets for immunotherapy. In one approach we used a set of peptides derived from Gag and Pol that were selected on the basis of extreme sequence conservation and coverage of a wide range of HLA associations, and in the other approach we used a set of topologically important Gag, Pol, Env, and Nef associated epitopes selected based on structural analysis. As a more conventional approach, we also tested a pool of overlapping peptides spanning the Gag proteome, which consisted of both conserved and highly variable epitopes from the Thailand strain CRF01. These three peptide pools were part of an MDC1-based vaccine strategy and tested for their capacity to stimulate HIV-specific T cell responses *in vitro*. As we expected, peptide antigen alone was unable to induce the activation and expansion of the isolated T cell cultures, but instead required the presence of the MDC1. Moreover, we did not see enhancement of HIV-specific responses when T cells were expanded in the presence of MDC1 in the absence of peptide. Notably, MDC1 that were loaded with either of the three peptide pools were consistently capable of generating the activation and expansion of HIV-specific T cells. These results are consistent with the notion that MDC1 are highly specialized as potent stimulators of T cell activation through their capacity to professionally present antigen in the context of MHC molecules. This triggers TCR signaling (signal 1), that along with costimulatory molecules (signal 2), provide timely, important cytokines, including IL-12p70 (signal 3), which together are critical to support the further expansion and survival of effector T cells [6].

When testing the larger peptides that were used to initiate the DC:T cell cocultures in the effector cell readout assays, we found that a majority of the T cell responses activated were within the CD4^+^ T cell fraction. This finding was most pronounced when we assessed responses against the Epigraph peptide pool. We speculated that our short-term readout assays were not detecting HIV-specific CD8^+^ T cells that were selectively expanded using the MDC1-based approach. We reasoned that this was because the peptides were not of optimal size for their direct MHC-class I loading and presentation in the assays. Indeed, a comparison between the use of a 21mer peptide taken from the afferent stimulator peptide pool cultures versus a 9mer derived from the same 21mer determined that the smaller peptide clearly revealed a higher magnitude of CD8^+^ T cell responses during the IFNγ secretion readout assay. These results are in concordance with previous evidence that 15mer sequences are not ideal antigens for identifying and enumerating MHC class I-restricted CD8^+^ T cell responses [40]. However, we showed that MDC1 can efficiently process the longer exogenous peptides for cross-presentation to drive a broad range of MHC-class 1 restricted CTL responses. This is in accord with previous studies demonstrating benefit of exposure of DC with longer exogenous peptides and/or proteins rather than shorter peptides to achieve prolonged cross-presentation on MHC class I and efficient afferent induction of CTL immunity [47–50].

In order to more accurately evaluate the induced effector CD8^+^ T cell responses by IFNγ ELISpot, we assessed a group of smaller 9-13mer stimulator peptides that were derived from the larger peptide sequences used to initiate the cultures. To select these smaller peptides for testing, we defined a number of optimal MHC-class I epitopes derived from the larger 14-21mer sequences that also had broad HLA associations specific to those haplotypes present in our study cohort using the LANL and IEDB. This allowed us to identify a wider range of CD8^+^ T cell responses than would have otherwise been revealed. However, this dramatically increased the number of peptide epitopes to be tested, the number of cells needed to carry out the assays, and the overall difficulty of monitoring the effectiveness of the MDC1-based vaccine strategy. Moreover, by selecting only those peptides associated with the HLA haplotypes within our cohort for the readout assays, we understand that a number of unknown but relevant epitope responses could have been missed, and thus highlights a limitation of our study. This is supported by our finding of a number of unexpected responses to peptides with known associations with HLA haplotypes other than those of the particular study participant being tested, suggesting the discovery of novel MHC class I epitopes that was revealed as a result of the initial MDC1 stimulation strategy. Nevertheless, our study highlights the difficulties of accurately assessing or monitoring the responsiveness to a vaccine that has the potential of eliciting responses to such a broad range of antigenic targets, with a large number of HLA-associations, through the use of standard monitoring techniques, and the need to be cognizant of these points when designing a vaccine study.

Detailed analysis of the data generated in this study led to additional noteworthy findings. One outcome was the observed higher magnitude of effector T cell response generated against several Network 9-13mer peptides, particularly in those individuals initiating ART in the Fiebig III stage of infection. In addition, a positive correlation was found between a study participant’s viral load at the initiation of ART and the magnitude of responses against the Network peptide pool. Although the sample size of individuals tested was too small to make definitive conclusions, we speculate that both the time to treatment and the antigen burden (viral load) at ART initiation had considerable impact on the capacity to induce primary CD8^+^ T cell responses against topologically important, conserved epitopes. As a result, this subsequently affects the quality of the anamnestic responses capable of being generated *in vitro*, which are what were measured here. These results are in accordance with a previous report showing that individuals in stage 3 (equivalent to Fiebig III) undergo full T cell differentiation during AHI and are therefore able to respond more effectively than individuals initiating ART before or after this window of peak viremia [51].

While the two methods employed to select the antigenic targets were strategically different, we hypothesized that both would share a high degree of conservation within the HIV-1 M group, and in particular within the CRF01 clade most relevant to the Thailand cohort. However, when the sequences were analyzed for their degree of exact matching to viruses found among the entire M group and those specific to the CRF01 clade, the Network peptides were not as relatively conserved as the Epigraph peptides (S4 Fig and S5 Fig).

However, this apparent variability within the Network epitopes was observed in peptides which were selected at an interim stage of development of the structure-based network analysis approach, and were in fact, not classified as being topologically important upon finalization of the algorithm (9). The modest polymorphism observed for topologically important epitopes that precludes them being classified as exact matches in HIV-1 M group is likely the result of their underlying immunogenicity, which we observed in this study. Moreover, mutations within highly networked epitopes have been shown to impair viral fitness, so these responses may be effective in mediating immune control despite mutational changes [9, 52–54]. The finding that the networked epitopes were targeted more robustly than the Epigraph peptides and show evidence of some sequence variation likely indicates that these are more immunogenic *in vivo*, and that these topologically important regions would indeed be represented as part of the peptide repertoire presented on the infected target cells. Moreover, while the extreme level of conservation among the Epigraph peptide sequences could mean that they are important for viral fitness and less likely to be subject of CTL immune escape, it could also indicate these peptides are not as readily presented on HIV-1 infected cells, and therefore may not be as reliable for immune targeting. Nevertheless, we believe that both antigen selection strategies for vaccine development have their distinct advantages, and therefore propose a mosaic model, where antigenic peptides representing both highly conserved and topologically important viral sequences are targeted.

Our study demonstrates that by using MDC1, it is possible to induce the activation and expansion of HIV-1 specific T cells capable of responding to highly conserved and topologically important epitopes in people initiating ART during early stages of infection. We demonstrated that autologous MDC1 can be loaded with larger exogenous peptide antigen that is cross-presented in the context of MHC class I as well as MHC class II, and used as a cellular vaccine platform to induce the activation and long-term expansion of both HIV-specific CD8^+^ T cells and CD4^+^ T cells, respectively. Most importantly, rather than targeting the viral proteome as a whole, by carefully choosing the target antigen to include only those regions of the virus that contain sub-dominant, ultra-conserved, and topologically important epitopes, the competition for MHC binding and presentation that may otherwise favor memory T cell responses toward highly variable immunodominant epitopes can be limited. By prospectively avoiding such ‘immunologic noise’, we demonstrate a level of ‘immune focusing’, i.e., where HIV-1 antigen-loaded MDC1 have a therapeutic competence to selectively drive and focus CTL responses toward highly conserved epitopes that are less likely to lead to viral escape from CTL pressure and more likely to prove critical to viral fitness [9]. Together, this study highlights the potential of implementing this MDC1-based approach for selective immune targeting as an integral part of a successful ‘kick and kill’ strategy to control chronic HIV infection.

## Materials and Methods

### Study cohort participants

The RV254/SEARCH 010 (NCT00796146 and NCT00796263) cohort enrolls adults diagnosed with acute HIV infection (AHI) at the time of presentation at an HIV screening site at the Thai Red Cross Anonymous Clinic and who were offered immediate ART [36, 55, 56]. The Chulalongkorn University Institutional Review Board and the Walter Reed Army Institute of Research, USA, approved this study. AHI is defined as either non-reactive 4^th^ generation immunoassay with positive nucleic acid test or reactive 4^th^ generation immunoassay together with non-reactive 2^nd^ generation immunoassay [57]. The procedures of staging AHI have been described previously [27, 36, 57]. For this study, peripheral blood mononuclear cells (PBMC) were obtained from 10 HIV-infected individuals in the Thailand/MHRP RV254 cohort who initiated ART during acute/early infection [Fiebig I (n=2), II (n=2), III (n=4), IV (n=2); See Table 1].

### Human leukocyte antigen (HLA) genotyping

HLA genotyping was performed using a multi-locus individual tagging-next generation sequencing (MIT-NGS) method as described previously [58]. Briefly, DNA was extracted from PBMC and full-length HLA genes were sequenced by NGS on the MiSeq platform (Illumina, San Diego, CA). FASTQ files generated by MiSeq Reporter were analyzed by NGSengine v2.16.2 (GenDX, Utrecht, The Netherlands).

### 14-21mer peptides used to initiate for T cell cultures

Conserved antigens were identified by (1) selecting regions spanning only the most conserved potential T cell epitopes in the HIV proteome, based on the Los Alamos HIV database, using the Epigraph tool [26], While some highly conserved regions were identified in Env and Nef, we did not include them in this study as our original intent was to focus on the 5’ side of the genome to facilitate sequencing of clinical samples in future studies. We lifted this constraint for (2) structure-based network analysis using graph theory metrics to identify structurally and functionally conserved epitopes [9] (Fig 1). Overlapping peptides spanning the entire Gag proteome were used as control antigens (Gag peptide pool). The library of peptides was synthesized by Sigma-Aldrich (St. Louis, MO) and each peptide was resuspended at a concentration of 5 mg/ml using either DMSO (for peptides with negative polarity) or DI water (for peptides with positive polarity). Resuspended peptides were aliquoted and stored at −80°C until use.

### Selection of 9-13mer epitopes within larger 14-21mer peptides for use as readout stimulating antigen

For selection of 9-13mer epitope sequence deriving from the larger afferent 14-21mer sequences (Gag, Network, and Epigraph), we identified known and predicted CD8^+^ T cell epitopes and their HLA associations, using the Immune Epitope Database (IEDB) and the Los Alamos National Laboratory Database (LANL) based on MHC class I binding predictions (IC50<500). We then selected those epitopes contained within each 14-21mer that were predicted to provide maximum coverage of different HLA-types represented in our group of study participants as our readout antigens.

### Isolation of peripheral blood monocytes and lymphocytes

PBMC from study participants were collected, aliquoted and frozen at a concentration of 40×10^6^ PBMC per vial. Cells were shipped to our facility and kept in liquid nitrogen until use. PBMC were thawed and monocytes and peripheral blood lymphocytes (PBL) were separated using human anti-CD14 Ab-coated microbeads to positively select monocytes (Miltenyi Biotec, Auburn, CA) according to the manufacturer’s instructions. Negatively isolated PBL were cryopreserved for future use.

### Generation of human monocyte-derived DC (MDC1)

Isolated monocytes were cultured for 5 days in Iscove’s Modified Dulbecco’s Media (IMDM) containing 10% fetal bovine serum and 0.5% gentamicin in the presence of 1000 IU/ml of human granulocyte-macrophage colony-stimulating factor (GM-CSF; R&D Systems, Minneapolis, MN) and 1000 IU/ml of interleukin 4 (IL-4; R&D Systems, Minneapolis, MN) to differentiate them into immature dendritic cells (iDC) in a 24-well plate. On day 5, iDC were divided into 4 groups of treatment, i.e., untreated (Empty; no peptide) or loaded with the 14-21mer Gag-overlapping peptide pool (Gag, n=45), the Network peptide pool (Network, n=25), or the Epigraph peptide pool (Epigraph, n=40), at a final concentration of 1μg/ml for each peptide. After a 2 h incubation at 37°C, a type-1 polarizing (MDC1) cytokine cocktail containing interleukin-1β (IL-1β; 25 ng/ml), tumor necrosis factor α (TNFα; 50 ng/ml), IFNγ (3000 IU/ml) (R&D systems), IFNα (1000 IU/ml) (Miltenyi Biotec) and poly (I:C) (20 μg/ml) (Sigma-Aldrich) was added to the iDC cultures for 48h to yield mature MDC1 as previously described [59, 60]. Once matured, MDC1 were harvested and exposed again to the 14-21mer peptide pools for 2 h prior to being used for T cell stimulation.

### *In vitro* stimulation of bulk T cells

MDC1 from the 4 different groups described above (Empty, Gag, Network and Epigraph) were counted and plated at a concentration of 7.5×10^4^ MDC1 per well in a 24-well plate. Bulk T cells were negatively selected using the EasySep^TM^ Human T Cell Enrichment Kit (STEMCELL Technologies, Cambridge, MA), and 7.5 x 10^5^ T cells were added per well to the MDC1 containing wells (MDC1 to T cell ratio = 1:10). After an incubation of 30-45 min at 37°C, soluble recombinant human CD40L was added at a concentration of 0.25 μg/ml (MEGACD40L^®^ protein, ENZO Life Sciences, Farmingdale, NY). After 4 to 5 d stimulation, rhIL-2 (250 IU/ml) and rhIL-7 (10 ng/ml) were added to the cultures and every 3 d thereafter. After a total of 21 d in culture, T cell responses against 9-13aa peptide epitopes derived from the longer (14-21aa) stimulator peptides were determined by IFNγ ELISpot and by CD107a flow cytometry staining.

### Surface and intracellular staining and flow cytometry

Expanded T cells were harvested after 21 d in culture, counted and plated in a V-bottom 96 well plate at a concentration of 1×10^5^ cells per well and rested overnight before stimulation with 9-13mer peptide pools. Antigen-specific T cell responses were assessed by CD107a staining and IFNγ intracellular cytokine staining. Cells were resuspended in media containing CD107a-FITC labeled antibody (Clone H4A3, BD Bioscience, San Jose, CA) and BD GolgiStop^TM^ (protein transport inhibitor containing monensin, BD Bioscience) according to the manufacturer’s instructions. Peptide pools containing 9-13mer peptide sequences were added to respective wells and incubated for 6 h at 37°C. Wells without peptide addition were used as controls. After incubation, cells were washed with 1X PBS and stained for viability with LIVE/DEAD^TM^ Fixable Aqua Dead Cell Stain Kit (Invitrogen^TM^ Molecular Probes^TM^) for 20 min at room temperature in the dark. Surface staining was done subsequently using anti-CD3, -CD4 and -CD8 antibodies and incubated for 30 min at room temperature in FACS buffer. After surface staining, cells were washed, fix and permeabilized using the BD Cytofix/Cytoperm^TM^ Fixation/Permeabilization Kit (BD Bioscience) and stained with IFNγ monoclonal antibody (IFNγ-AlexaFluor® 700, clone B27; BD Bioscience) for 45 min in the dark. Samples were acquired in a LSR Fortessa II (BD Bioscience) flow cytometer and subsequently analyzed using the FlowJo software (Tree Star).

### ELISpot for detecting IFNγ secreting cells

*In vitro* expanded T cells were harvested, counted and immediately used for IFNγ secretion by ELISpot. The IFNγ ELISpot assay was performed following the Mabtech Human IFNγ ELISpot^Basic^ protocol (Mabtech, Cincinnati, OH) using 96-well PVDF ELISpot plates from Millipore, as previously described [21]. Briefly, T cells were resuspended at a concentration of 3×10^5^ cells per milliliter and 100 μl (3×10^4^ cells/well) were transferred to IFNγ-antibody coated 96-well ELISpot plates (5μg/ml anti-IFNγ mAb 1-D1K, Mabtech, Stockholm, Sweden). Individual 9mer peptide dilutions were done in a separate 1 ml deep-well 96-well plate at a final concentration of 2μg/ml and 100 μl of this dilution was added to T cell-containing wells to give a final peptide concentration of 1μg/ml. All ELISpot assays included negative-control wells with expanded T cells without peptide stimulation (Media only). T cells expanded using control MDC1 without peptide were also tested for responses to the respective 9-13mer peptide pools but yielded no antigen-specific responses (data not shown). IFNγ responses to each peptide were done in duplicate wells. The enumeration of spots was done using the Autoimmun Diagnostika GmbH (AID) EliSpot reader and counting software (AID, Strassberg, Germany). ELISpot data were calculated as the means of spots in duplicate wells minus the mean and 2 standard deviations of the negative control values and shown as IFNγ spots/ 10^6^ cells. We defined a positive responder as having a calculated ELISpot value larger than 50 SFU/10^6^ cells.

### Statistical analysis

The statistical analyses and plotting of data were performed using GraphPad Prism 8 software. A non-parametric method, signed-rank test was used to evaluate the equality of matched pairs of observations by using Wilcoxon matched-pairs for paired data and single sample signed test for the normalized data. We used a two-sided P value < 0.05 to highlight differences of interest; note that multiple tests were performed, and these are uncorrected p-values, thus these values should be considered as indicative of trends of interest in the data.

## Acknowledgements

This study was supported by the NIH, National Institute of Allergy and Infectious Diseases grants R21-AI131763, R21-AI138716, UM1-AI126603, and U01-AI35041. We thank Holly A. Bilben and Jan Kristoff for technical support. We would like to thank the RV254/SEARCH 010 study participants and support staff who committed their time and effort for this study. The RV254/SEARCH 010 is supported by cooperative agreements (WW81XWH-18-2-0040) between the Henry M. Jackson Foundation for the Advancement of Military Medicine, Inc., and the U.S. Department of Defense (DOD) and by an intramural grant from the Thai Red Cross AIDS Research Centre. This research was funded, in part, by the U.S. National Institute of Allergy and Infectious Diseases. Antiretroviral therapy for RV254/SEARCH 010 participants was supported by the Thai Government Pharmaceutical Organization, Gilead, Merck and ViiV Healthcare.

## DISCLAIMER

The views expressed are those of the authors and should not be construed to represent the positions of the U.S. Army or the Department of Defense. The investigators have adhered to the policies for protection of human subjects as prescribed in AR 70–25.

