## Supplemental Figures for "Dendritic cells focus CTL responses toward highly conserved and topologically important HIV epitopes"

**Garcia-Bates, S1 Fig**

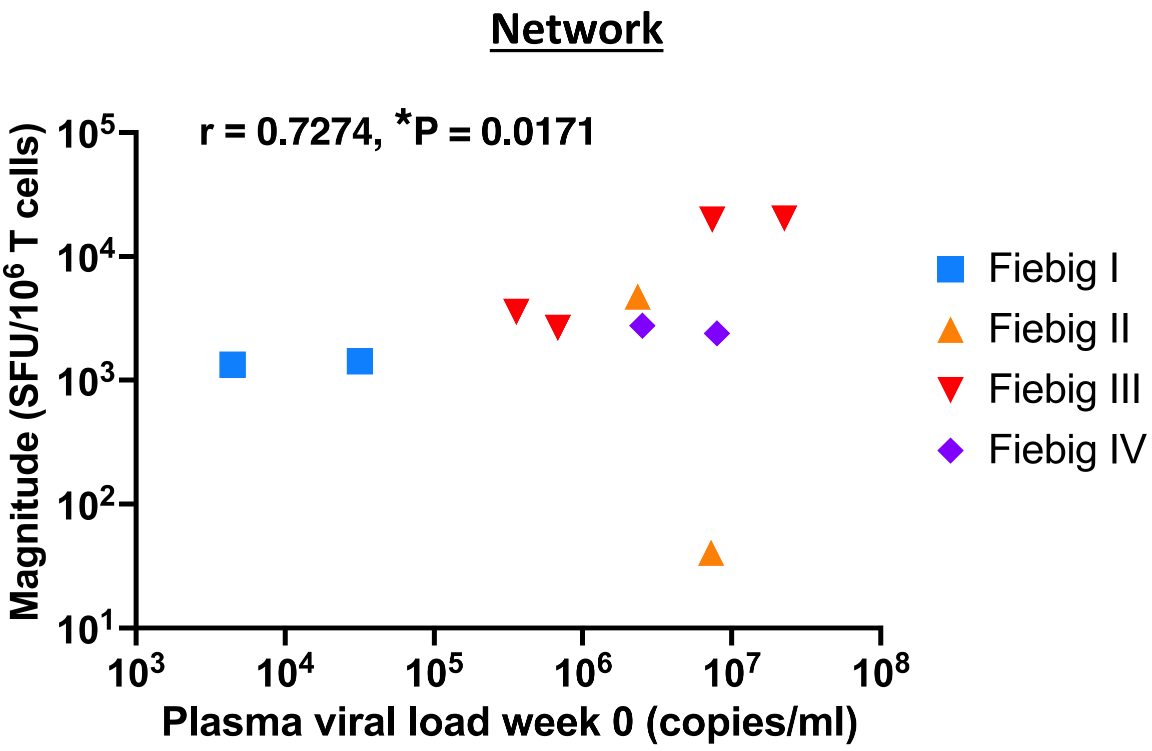


**S1 Fig.** Correlation between plasma viral load at week 0 prior to ART treatment and magnitude (SFU/$10^6$ T cells) of IFN$\gamma$ responses for each Fiebig stage for AHI.

### Garcia-Bates, S2 Fig

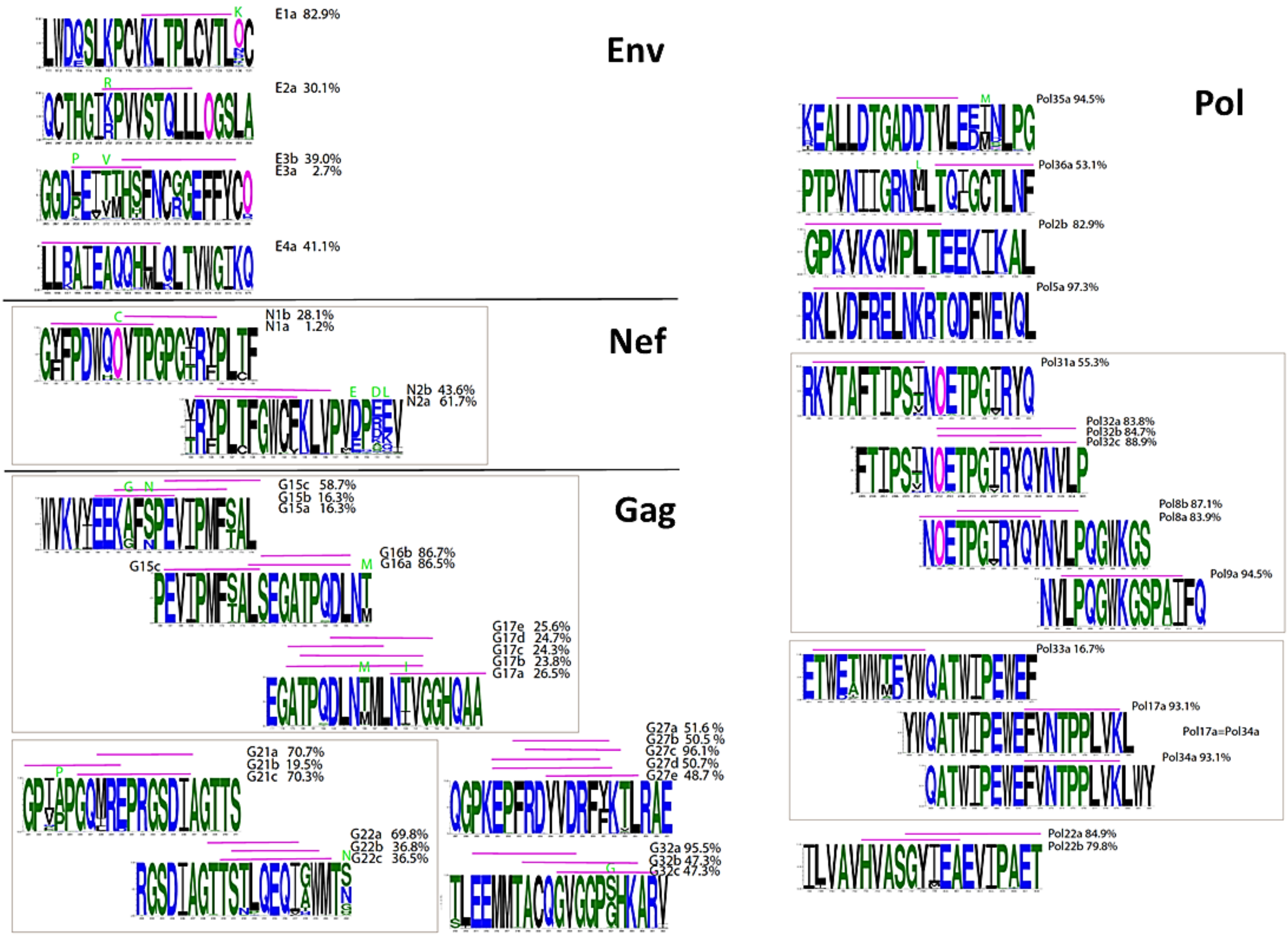


**S2 Fig. Network peptide variability for Env, Nef, Gag and Pol proteins.** Logos show the frequency of the amino acid in the **M group** in the network peptides. The most common amino acid is on the top. Magenta O's are N's embedded in glycosylation sequences, Nx[ST]. Green letters above the logo means the amino acid within network peptide does not match the most common form globally. Pink lines are the efferent epitopes for the respective afferent peptide. The number following the designation, is how many times an exact match is found in the M group.

### Garcia-Bates, S3 Fig

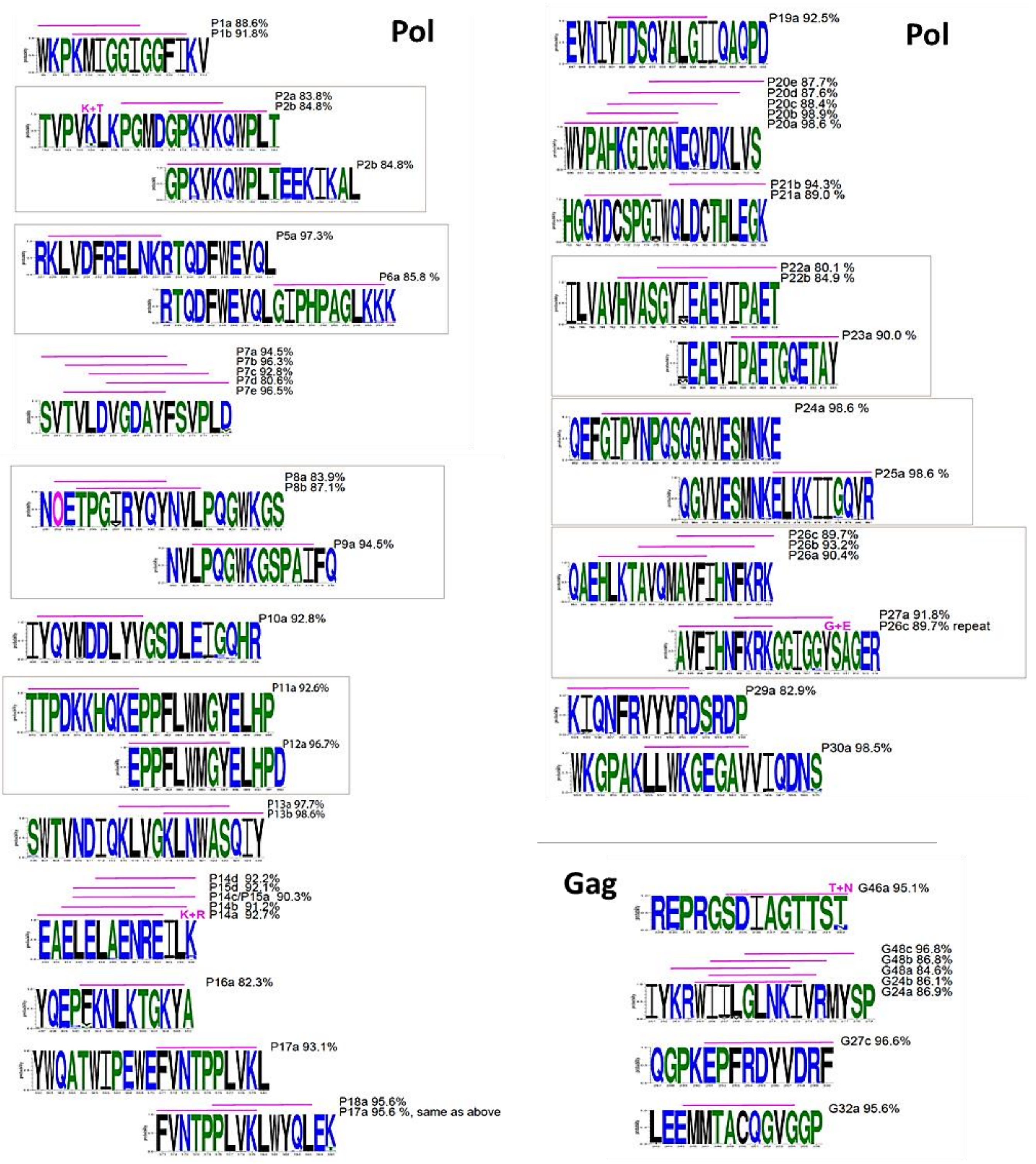


**S3 Fig. Epigraph peptide variability for Pol and Gag proteins.** Logos show the frequency of the amino acid in the **M group** in the network peptides. The most common amino acid is on the top. Magenta O's are N's embedded in glycosylation sequences, Nx[ST]. Green letters above the logo means the amino acid within network peptide does not match the most common form globally. Pink lines are the efferent epitopes for the respective afferent peptide. The number following the designation, is how many times an exact match is found in the M group.

**Garcia-Bates, S4 Fig**

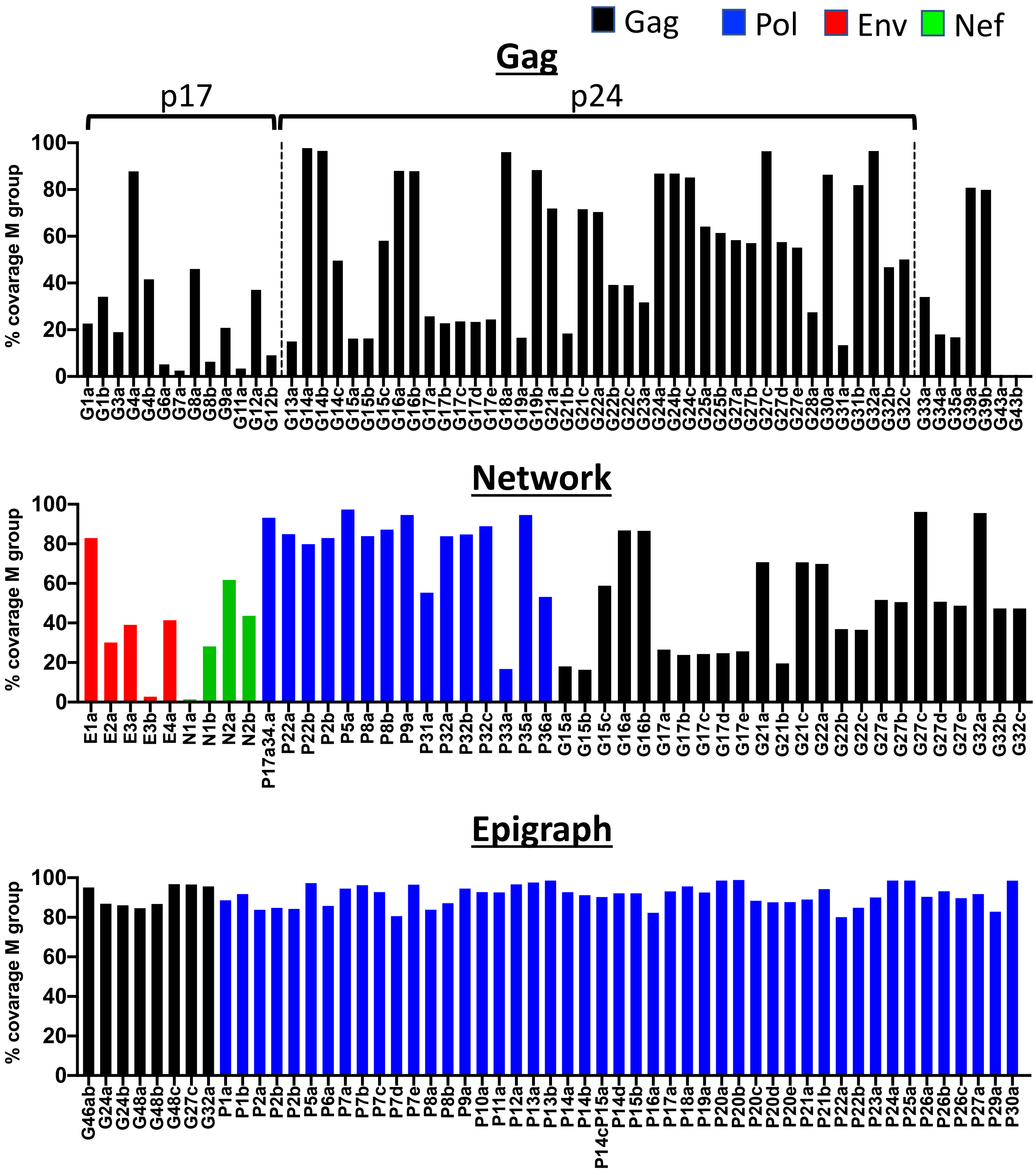


**S4 Fig. Exact matching frequency of epitopes with the M group alignment**

Garcia-Bates, S5 Fig

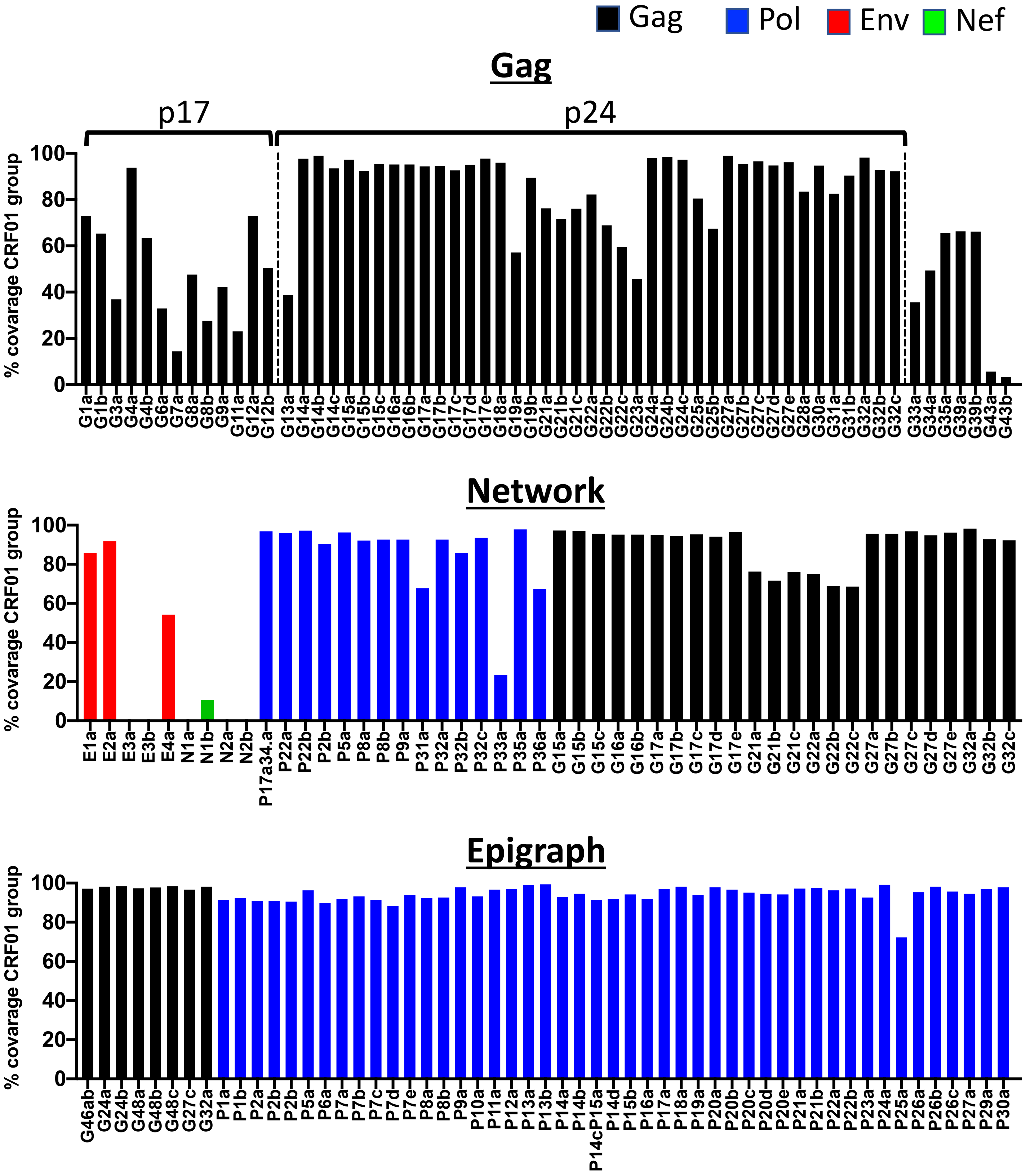


S5 Fig. Exact matching frequency of epitopes with the CRF01 group alignment
